# The landscape of autosomal-recessive pathogenic variants in European populations reveals phenotype-specific effects

**DOI:** 10.1101/2020.11.16.384206

**Authors:** Hila Fridman, Helger G. Yntema, Reedik Mägi, Reidar Andreson, Andres Metspalu, Massimo Mezzavila, Chris Tyler-Smith, Yali Xue, Shai Carmi, Ephrat Levy-Lahad, Christian Gilissen, Han G. Brunner

**Author notes:** These authors contributed equally (Ephrat Levy-Lahad, Christian Gilissen, Han G. Brunner).

## Abstract

The number and distribution of recessive alleles in the population for various diseases are not known at genome-wide-scale. Based on 6447 exome-sequences of healthy, genetically-unrelated Europeans of two distinct ancestries, we estimate that every individual is a carrier of at least 2 pathogenic variants in currently known autosomal recessive (AR) genes, and that 0.8-1% of European couples are at-risk of having a child affected with a severe AR genetic disorder. This risk is 16.5-fold higher for first cousins, but is significantly more increased for skeletal disorders and intellectual disabilities due to their distinct genetic architecture.

---

A major public health goal is to detect at-risk couples (ARCs) for various autosomal-recessive (AR) diseases. Detecting such couples would enable them to consider reproductive choices to prevent the birth of affected children, including prenatal diagnosis (PND) and preimplantation genetic testing (PGT). Currently, the number and distribution of recessive alleles in the population is not known at genome-wide scale. Understanding the architecture of AR pathogenic variants can contribute to the knowledge base for public health policies in the preconception field, and illuminate the evolution of disorders and phenotypes.

Existing estimates of the number of AR alleles carried by individuals are either derived from comparisons of the incidence of AR disorders between offspring of consanguineous and non-consanguineous couples, or based on extrapolations from sequencing data of specific phenotypes and gene-sets. Early calculations estimated that each individual carries at least eight heterozygous recessive pathogenic variants^1^, while estimates based on consanguineous couples predicted 3-5 heterozygous recessive lethal pathogenic variants per individual^2^. Later models predicted up to 100 pathogenic variants per individual^3^. An analysis based on gene-dropping simulations in a well-documented isolated founder population (The Hutterites), estimated that each founder of this population carried 0.58 AR recessive lethal pathogenic variants that lead to death between birth and reproductive age or to complete sterility^4^.

Sequencing-based studies of wider gene-sets also yielded variable results. Screening for 437 known AR genes related to Mendelian diseases found a carrier frequency of 2.8 severe pathogenic variants per individual (range 0-7)^5^. In another study that used samples of various ethnicities including Caucasians, testing for a panel of 417 AR pathogenic variants found ~0.4 AR lethal pathogenic variants per individual, leading the authors to suggest that the number for the entire genome is ~10 times higher^6^.

None of the existing studies used direct gene-sequencing at a genome-wide scale. Furthermore, each study used a different methodology, cohort size and number of tested genes and variants. Therefore, current data do not allow an overall assessment of the genomic landscape of AR disease variants.

Here, we performed an exome-sequencing based assessment of the carrier frequency of AR pathogenic/likely-pathogenic variants (PLPs), the total ARCs rate for various disorders, and the effect of different consanguinity levels on the ARCs rate for these disorders in two distinct European populations. We used direct gene-sequencing and included a comprehensive set of AR genes. These analyses reveal the architecture and distribution of AR pathogenic variants throughout the genome and for different disorders. The results can inform public health policies such as design of preconception carrier screens and improve preconception counseling. Our results also provide novel insights into the population genetics of AR disorders, particularly regarding intellectual disabilities.

## Results

We gathered exome samples from two cohorts of healthy individuals of Dutch (n=4120) and Estonian (n=2327) populations, and filtered these for quality, kinship and ethnicity (**Fig. 1**). For these cohorts we analyzed a set of 1929 AR disease genes (**Methods; Supplementary Table 1**), including a subset of 1119 genes that were previously categorized as associated with severe phenotypes by manual expert curation as part of the development of an Australian pre-conception screening (PCS) panel^7^ (**Methods**). In these genes, we selected all pathogenic and likely pathogenic variants (PLPs) based on existing classifications from databases and ACMG guidelines^8^ (**Methods**). The filtering process was applied to a total of 91,341 and 45,929 variants for the Dutch and Estonian cohorts, respectively, and excluded >95% of the variants as either benign/likely benign or variants of unknown significance (VUS), resulting in 3734 and 1664 PLPs for the Dutch and Estonian cohorts, respectively (**Fig. 1, Supplementary Fig. 1, Methods**).

**Figure 1.**
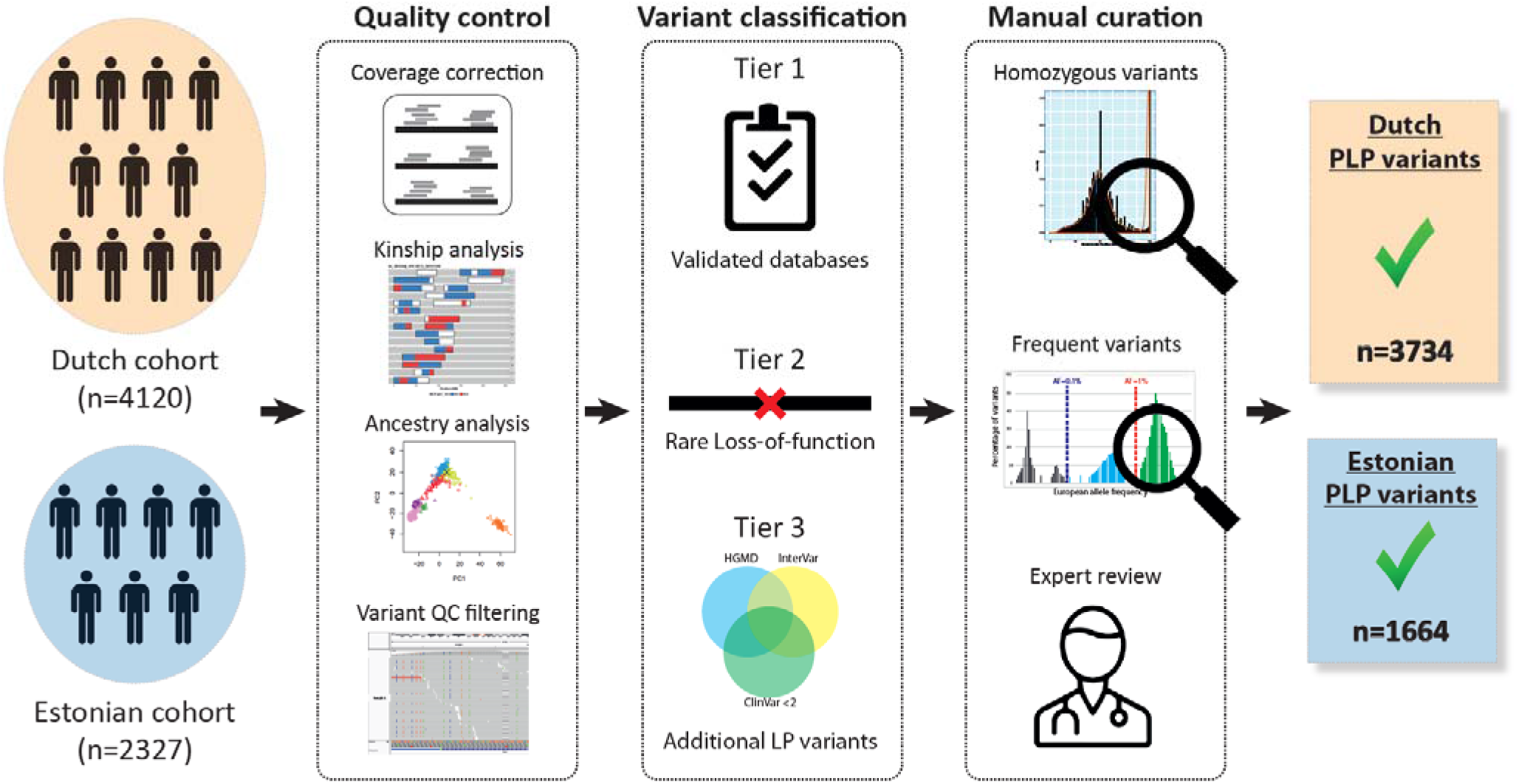
Overview of the selection of PLP variants. From left to right, variants were selected from two exome-sequencing cohorts of healthy individuals from two different European populations. **Quality control:** Samples and variants were subjected to stringent quality control. Samples were filtered for kinship and ethnicity. Variants were filtered for quality and selected from consistently well-covered regions. **Variant classification:** Variants were classified as PLP based on curated publicly available databases and/or on their predicted loss of function effect. **Manual curation:** We performed manual curation steps at both the gene and the variant level, to confirm the validity of our PLP classification selection process. Detailed information is in the **Methods** and **Supplementary Fig. 1**.

### Pathogenic variants

More than half of the PLPs (55.2%/59.1% in the Dutch/Estonian cohort) are rare Loss-of-Function (LoF) variants not previously described in Clinvar^9^ or the Dutch society of laboratory specialists initiative for data sharing of variant classifications (VKGL database^10^; www.molgenis.org/vkgl) (classified as PLP by tier 2 criteria) (**Supplementary Fig. 1**). About a third are known PLPs in the VKGL^10^ database and/or classified as PLP by ClinVar^9^ with a status review of 2 or more stars (34.6%/27.6% in the Dutch/Estonian cohort) (tier 1 criteria). The remaining 10.1%/13.2% are variants classified as PLP by ≥2/3 databases (tier 3 criteria) (**Methods, Supplementary Fig. 1**).

Since the classification of variants as PLPs forms the basis of this study we performed several analyses that confirmed the validity of our classification methodology (See **Methods, Supplementary Fig. 1**).

When comparing the two cohorts, we found that only 373 PLPs (10% of the Dutch PLPs and 22% of the Estonian PLPs) were shared between both cohorts (**Fig. 2a**). The 90% (n=3361) of unique Dutch PLPs had an average allele frequency of 3·10^-6^, and the 78% (n=1291) of unique Estonian PLPs had an average allele frequency of 2·10^-6^ (**Fig. 2a**). This highlights the difference in the underlying genetic architecture between the two populations and suggests that results obtained for each of these cohorts are largely independent, in that they are not based on the same genetic variation. This was to be expected since the Dutch and Estonian populations are separated geographically with limited interaction over recent history^11^ (**Supplementary Fig. 2**).

**Figure 2.**
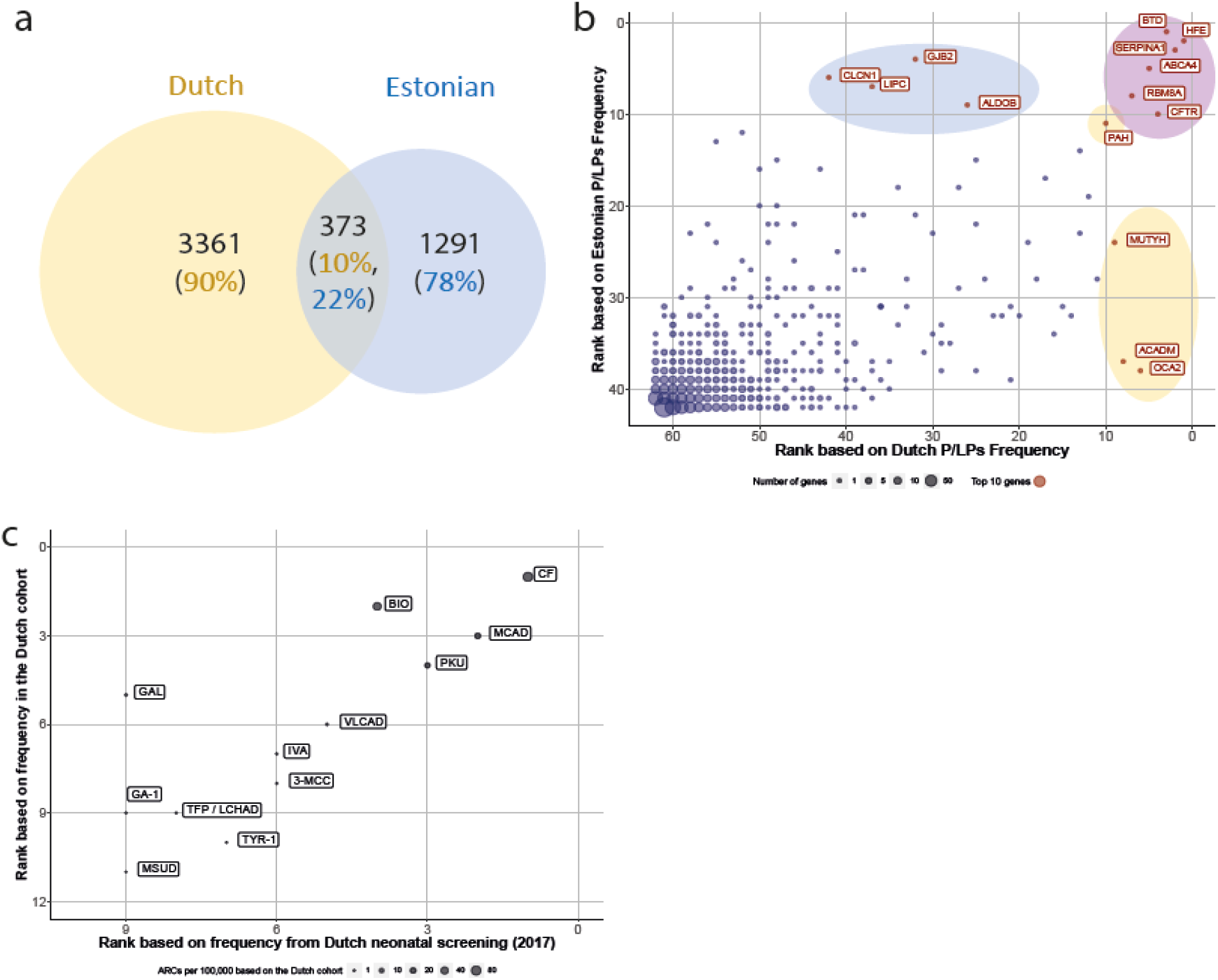
PLP variants in the Dutch and Estonian cohorts-robustness of variant classification. **(a)** <u>The number of PLPs in each cohort and their overlap</u>. **(b)** <u>Genes ranked by PLPs frequency: correlation between the Dutch and the Estonian cohorts</u>. The lower the rank number, the higher the number of PLPs observed in that gene (i.e., a gene ranked 1 has the largest number of PLPs); Genes with no PLPs in both cohorts were excluded (449 genes); Gene names are written for genes in the top 10 ranking: blue-Estonian only top-10, orange-Dutch only top-10, purple-Dutch and Estonian top-10; The dots sizes represent the number of genes with this rankings combination. Spearman correlation coefficient: 0.69; P-value<.00001. **(c)** <u>Comparison of disease carrier frequency estimates to published data of the Dutch neonatal screening program from 2016-2017</u> (www.rivm.nl/documenten/monitor-van-neonatale-hielprikscreening-2016;www.rivm.nl/documenten/monitor-van-neonatale-hielprikscreening-2018). Spearman correlation coefficient 0.85; P-value=0.0005; Full diseases and gene names are in **Supplementary Table 11**.

### Carrier frequency in the European population

On average, each individual carries 2.3 (range 0-11)/2.0 (range (0-9) PLPs for the set of 1929 AR genes, in the Dutch/Estonian cohort respectively (median 2/2; **Table 1**). For the subset of 1119 recessive genes that are associated with severe phenotypes, the mean number of PLPs per individual is 1.5 (range 0-8)/1.1 (range 0-6) in the Dutch/Estonian cohort (median 1/1; **Table 1**). In the Dutch/Estonian cohort, there were 397 (19.6%)/315 (13.5%) individuals with no PLPs, and 144 (3.5%)/29 (1.2%) individuals with more than 5 PLPs (**Supplementary Fig. 2**). Overall, our results establish that on average, Europeans carry at least 1 PLP variant for a severe AR disorder and ≈2 PLPs for any AR disorder.

**Table 1.** PLP variants and ARCs for AR genes in the study cohorts

|  |  | **Mean/median**<br>**P/LPs per**<br>**sample** | **ARCs %(N)**<br>**(Dutch total: 8,485,140**<br>**Estonian total: 2,706,301)** |
| --- | --- | --- | --- |
| **1929 genes** | **Dutch cohort**<br>**(N=4120)** | 2.3/2 | 1.5% (124,722) |
|  | **Estonian**<br>**cohort**<br>**(N=2327)** | 2/2 | 1.3% (34,570) |
| **1119 severe**<br>**genes** | **Dutch cohort**<br>**(N=4120)** | 1.5/1 | 1% (83,878) |
|  | **Estonian**<br>**cohort**<br>**(N=2327)** | 1.1/1 | 0.8% (20,710) |

### Frequency of PLPs per gene

Having established the number of PLPs that an individual carries for recessive disorders, we wanted to investigate which genes have the highest carrier frequencies and have the largest effect on ARCs rates. Most variants in both cohorts are rare, with a carrier frequency of up to 0.05% (**Supplementary Fig. 3**). As a result, 96.6% of the 1929 genes had a total PLP carrier frequency of no more than 0.5% in both cohorts (**Supplementary Fig. 4**). Of these, 589 (30.5%)/1012 (52.5%) were genes for which PLP carriers were not observed. At the other end of the distribution, there were 30 (1.6%)/24 (1.3%) genes with more than 1% PLP carrier frequency in the Dutch/Estonian cohort (**Supplementary Fig. 4**). There is an overlap between common genes in both populations, as 23 genes have more than 0.5% carrier frequency in both populations (**Supplementary Table 2**).

Ranking genes by the frequency of PLP carriers demonstrated good correlation between the two populations (Spearman’s correlation coefficient Rho=0.69, P-value=5.05·10^-276^). The exceptions were 8 genes in which recurrent PLPs are very common in one population and not in the other (**Fig. 2b**). Six genes were shared in the top-10 rankings of both cohorts (P-value=0.0001, permutation test; **Methods**). In conclusion, although the cohorts are independent and each cohort has its own unique variants, the patterns of gene carrier frequencies are similar for both cohorts (**Fig. 2b**).

To validate our per-gene carrier frequency estimates, we compared our results to the published 2016-2017 data of the Dutch neonatal screening program (www.rivm.nl/documenten/monitor-van-neonatale-hielprikscreening-2016;www.rivm.nl/documenten/monitor-van-neonatale-hielprikscreening-2018). We compared the rankings based on the observed frequency of the tested disorders to our data-based estimates and found an essentially complete correlation (**Fig. 2c**, Spearman’s correlation coefficient Rho=0.99, **Supplementary Table 3**).

### ARCs in the European population

To determine the rate of ARCs, we simulated all possible virtual matings among the Dutch cohort (n=8,485,140 matings) and among the Estonian cohort (n=2,706,301 matings; **Methods**). Simulations for all 1929 AR genes resulted in 124,722 (1.5%)/34,570 (1.3%) ARCs in the Dutch/Estonian cohort (**Table 1**), representing virtual matings in which both partners carried a PLP variant in the same gene. Couples in which both partners carried PLPs that are known to cause mild/asymptomatic phenotype in homozygotes were excluded from this analysis (**Supplementary Table 4**). Simulations of the subset of 1119 severe genes yielded 83,878 (1%)/20,710 (0.8%) ARCs in the Dutch/Estonian cohort (**Table 1**). Therefore, we estimate that 0.8%-1% of European couples are at risk for a child with a severe AR condition, and at least 1.3%-1.5% of couples are at-risk for any AR condition. Considering all 1929 genes, 90% of the ARCs are explained by the 115/84 most frequent genes in the Dutch/Estonian cohort. For the 1119 genes that are associated with severe disease, 90% of the ARCs are explained by the 70/57 most frequent genes in the Dutch/Estonian cohort (**Supplementary Fig. 9; Supplementary Table 5**). Since most ARCs are explained by a limited number of genes, adding more genes to existing PCS panels for non-consanguineous couples is not expected to substantially increase the PCS yield, due to diminishing marginal returns.

### Effect of consanguinity on ARCs in the European population

Consanguineous unions are not common in European populations, but are common worldwide and are increasing in Europe due to immigration. Previously, consanguinity has been estimated to occur in ~0.06% of couples of Dutch descent^12^, similar to other European countries^13,14^. Consanguineous couples are typically the first to be referred for preconception screening, because of their increased risk for recessive disease. However, the precise magnitude of this increased risk is unclear, and it is unknown whether it is the same for different disorders.

We simulated consanguineous matings based on the Hardy-Weinberg principles and calculated the expected risks for different degrees of consanguinity, relying on the count of shared alleles that is expected by the relationship (**Methods**). We estimate that for any AR disorder, the rate of ARCs is 20.9%-24.9% for first-cousins, 10.4%-12.4% for first cousins once-removed and 5.2-6.2% for second cousins. The ARCs rate for third cousins is 1.3%-1.6%. i.e. not different from that of non-consanguineous unions (**Supplementary Table 6**).

For first-cousins unions, considering all 1929 genes, 90% of the ARCs are explained by the 749/540 most frequent genes in the Dutch/Estonian cohort (**Supplementary Table 5**). This shows that diminishing marginal returns effect is not seen in consanguineous couples, and therefore these couples are expected to derive a greater yield from an exome-based PCS, in comparison to non-consanguineous couples.

### ARCs per phenotype in consanguineous and non-consanguineous matings

We compared ARCs rates for consanguineous vs. non-consanguineous matings for all genes (1929 genes) and for the sub-group of severe genes (1119 genes). We also performed this comparison for gene-groups based on diagnostic gene panels corresponding to 12 different types of disorders (**Supplementary Table 1**).

We first investigated the allele-counts for PLPs per gene per panel. We found striking differences in the distribution of allele-counts between the different disorders (**Supplementary Fig. 5**). For example, only a small fraction of the genes for ID have high (>10) allele-counts (8%/1% in the Dutch/Estonian cohort), compared to other panels. For example, many more deafness genes have high allele-counts (24%/11% in the Dutch/Estonian cohort). Next, we calculated the expected number of ARCs per panel for first-cousins vs. non-consanguineous couples. For each disorder, the fold-increase in ARCs due to consanguinity (first-cousins) is indicated as the Consanguinity Ratio (CR). The CR was 16 for all genes combined, indicating a 16-fold higher risk for first-cousins than for unrelated couples across the entire dataset (**Fig. 3c**). The CR for different phenotypic groups of disorders is consistent between the two populations (Spearman correlation 0.5; rising to 0.75 when excluding two common Estonian variants in *CLCN1* and *GJB2*) (**Fig. 3d**). Notably, we find that the CR is significantly higher for ID and skeletal disorders compared to the average of all genes, in both cohorts (**Fig. 3, Supplementary Fig. 6, Supplementary Table 12**). Thus, while consanguinity generally elevates the risk for an affected child with all AR conditions, this elevated risk is not the same for different disorders (**Fig. 3c**).

**Figure 3.**
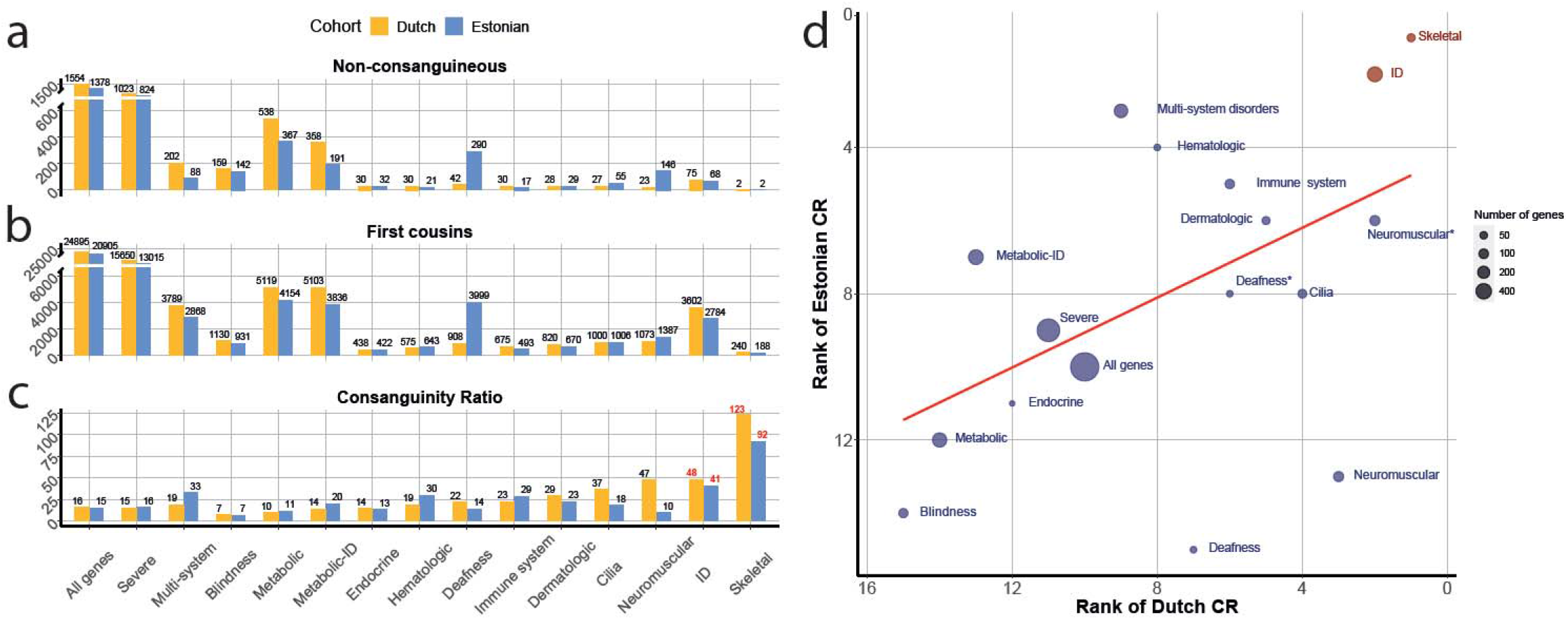
Effect of consanguinity on ARCs rates for different diseasecategories. **(a)** Rates of ARCs per 100,000 couples (on the y-axis) for different disorders (on the x-axis) in the Dutch cohort (orange) and Estonian cohort (blue) for non-consanguineous couples. **(b)** Same as in (a) but for first-cousin couples. **(c)** Consanguinity-ratio (CR) scores. Scores in red are significantly higher compared to a random set of AR genes with the same coding bp length. **(d)** Correlation of first-cousins CRs between the Dutch and Estonian cohorts for different disorders. X-axis shows the rank in the Dutch cohort; Y-axis shows the rank in the Estonian cohort. The lower the rank number, the higher the CR in that gene panel (i.e. a gene-panel ranked 1 has the highest CR). The size of the dots indicates the number of genes in the gene panel. The red color indicates gene panels with statistically significant higher CRs compared to a random set of AR genes with the same coding bp length. Rankings excluding common variants in *GJB2* (deafness) and *CLCN1* (neuromuscular) are marked with an asterisk.

Based on the allele-counts and CR scores, we calculated the expected distribution of disorders among affected children (**Fig. 4**). In the Dutch cohort, Metabolic disorders and blindness constitute 79% of expected disorders for affected children to non-consanguineous parents, while they only constitute 55% for affected children to parents who are first-cousins. Other phenotypes like ID and skeletal disorders are expected to be very rare in affected children to non-consanguineous parents, but much more common in children to parents who are first-cousins (**Fig. 4**).

**Figure 4.**
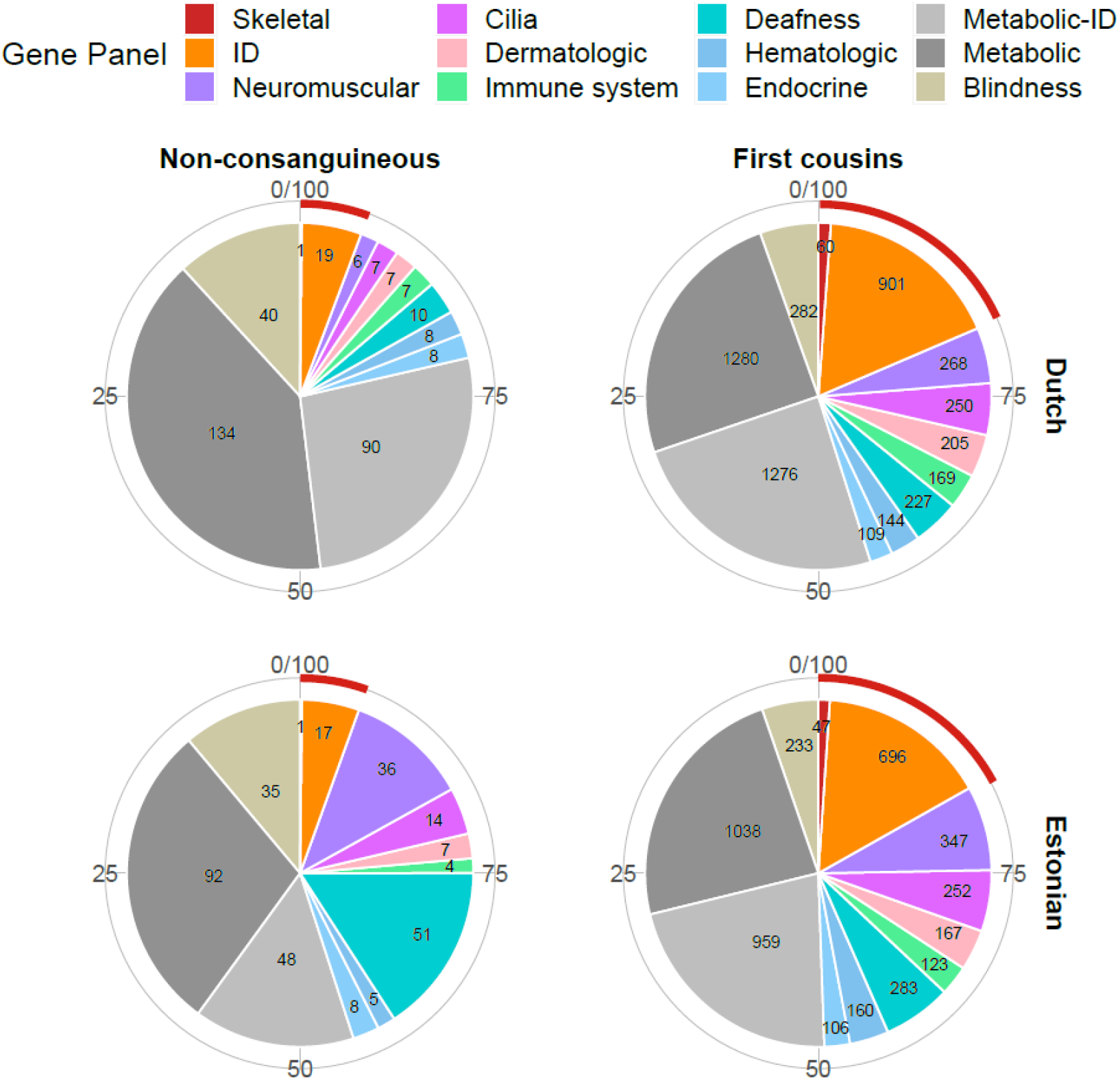
Effect of consanguinity on affected offspring rates for different disorders. The expected number of affected children for different disorders per 100,000 births, in non-consanguineous and first cousins couples. The red outer arch represents disorders with significantly higher CR scores. The difference between the cohorts for the metabolic gene panel is mostly attributed to a high carrier frequency in two genes (*CFTR* and *SERPINA1*) in the Dutch cohort.

### Heterozygote selection as a possible cause for the differences of PLPs patterns among disorders

A possible reason for the difference in the genetic architecture of ID and skeletal disorders compared to other disorders might be a fitness effect for heterozygous carriers of pathogenic variants in ID/skeletal genes. It is well-known that in some AR diseases there is indeed a phenotypic manifestation in heterozygotes^15,16^. Simulations show that even if heterozygosity for deleterious AR alleles reduces fitness only mildly, this would greatly reduce the frequency of variants in recessive genes for ID and skeletal disorders. In particular, if heterozygotes for a PLP variant have 0.5% less offspring (reduced fitness), in a large population this PLP variant will not have a frequency higher than 0.09% (**Supplementary Fig. 7**)^17^. Based on this hypothesis, we investigated the density of coding singleton variants (i.e. coding variants reported in only one individual) for the different gene panels in the 1000 Genomes dataset^18^ (**Fig. 5**). This dataset contains genome-sequences samples from 5 different European populations (GBR, TSI, FIN, IBS and CEU). We found that among these various European populations, the ID and skeletal disorders genes show a decreased number of coding singletons compared to the other gene sets, and are more similar to a set of essential genes that includes genes that are more likely to be under selection (**Fig. 5**). Similar patterns were observed for the singleton density across the entire gene, and for the RVIS (Residual Variation Intolerance Score) that is based on the number of functional variants in a gene (**Supplementary Fig. 8**). In conclusion, these results suggest that genes in the ID and skeletal panels are subject to increased selection pressures in the European population.

**Figure 5.**
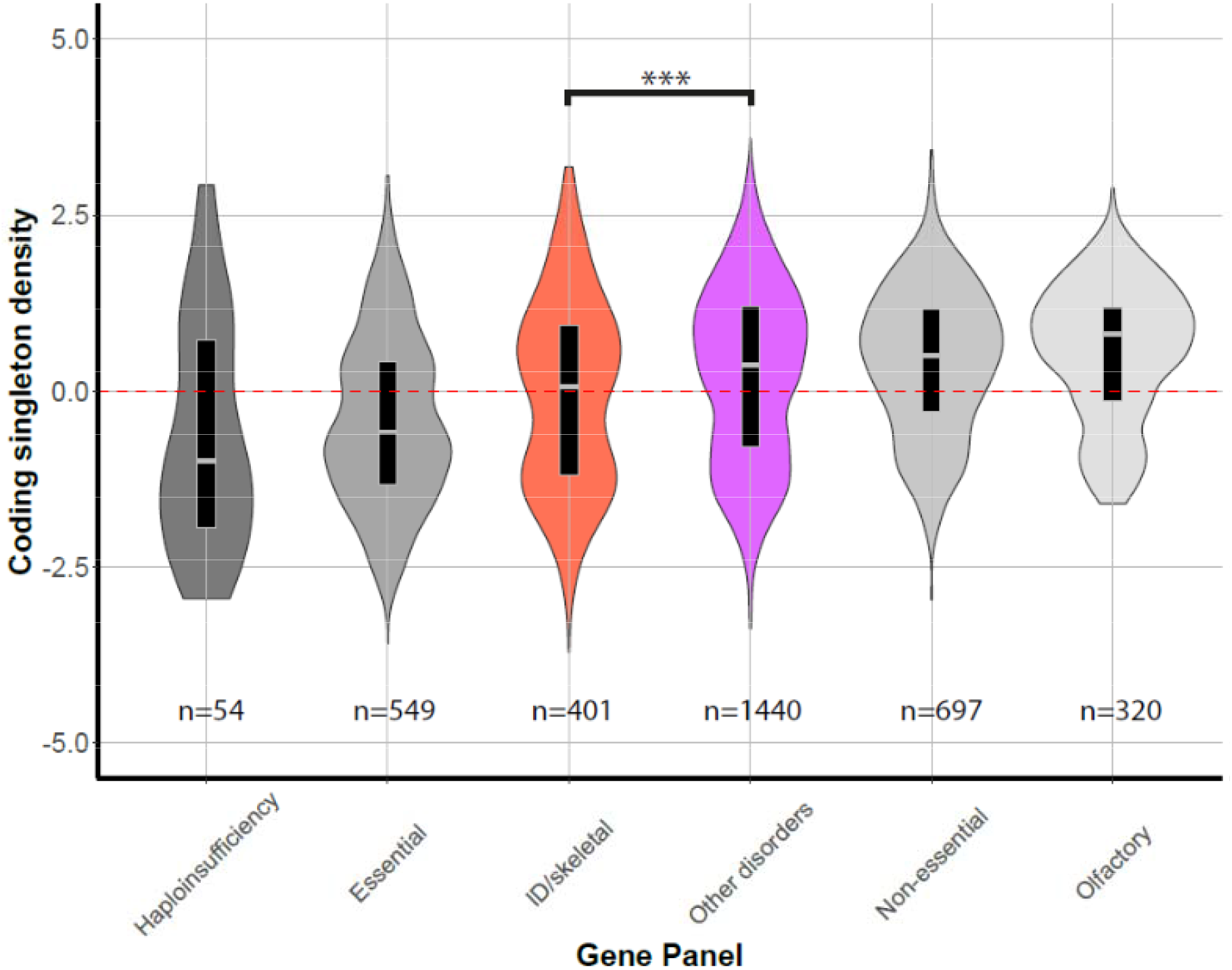
Coding singleton density in coding regions based on 1000 genomes data (European populations) for the different gene panels. Violin plots of the distribution of coding singleton density scores for different gene panels. Gray colors indicate external reference gene sets, whereas colored violin plots are gene sets as defined in this manuscript. Tested differences of the median are indicated by braces, giving rise to a P-value of 1.8·10^-4^ for ID/skeletal gene set compared to “other disorders” gene set, using a Wilcoxon rank sum test with Bonferroni correction (**Supplementary Table 13**).

## Discussion

We found that almost all individuals (>85%) carry at least one PLP variant, with an average of at least 1.3 PLPs for a severe AR disorder. Analysis of virtual matings for each population shows that in the absence of consanguinity, the rate of ARCs is 0.8-1% for a severe AR disorder. This translates to ~225 newborns with a severe AR disorder per 100,000 births. We believe these should be considered as minimal estimates. Firstly, we analyzed only the genes that are currently known as AR disease genes, while many new AR genes are still being discovered. Secondly, we took great care to avoid (likely) benign variants and VUS in our analysis, with the likely result of having excluded some variants that are actually pathogenic. This applies mainly to missense variants for which it is most challenging to predict their phenotypic effect and to hypomorphic variants. Lastly, in our analysis we did not consider variants in regions that are poorly covered by exome-sequencing, and other types of variation that are difficult to identify using exome-sequencing, such as intronic variants and copy number variation. Notably, the common SMN1 exon 7 and 8 deletion variant is not present in our data. Future analyses of whole genome sequencing data may give us the opportunity to obtain even more comprehensive estimates, although the systematic interpretation for these other types of variation will pose a significant challenge.

Crucial to our approach is the fact that we employed expert manual revision of classified variants. The current ACMG-based variant classification scheme is focused on pathogenicity, but does not consider the degree of pathogenic effect. Thus, two variants classified as “pathogenic” in the same gene, may have very different phenotypic effects. For example, in the *CFTR* gene, both deltaF508 and R117H are classified as pathogenic, but, whereas deltaF508 will result in classic cystic fibrosis, R117H may remain undetected or lead to mild disease. To avoid such problems, expert manual revision of classified variants cannot be spared from the classification process in individual cases.

Our results also underline the importance of population-specific databases. As seen in **Supplementary Table 7**, up to 30% of PLPs in 4 major gene-panels (deafness, blindness, ID, and metabolic disease), largely rare missense variants, were recognized only when we included information from the Dutch VKGL^10^ database. As expected, the number of PLPs added based on this part of the selection process was higher for the Dutch population than for the Estonian population (**Supplementary Table 8**). General worldwide databases currently do not include population-specific, unique, rare missense variants and thus, local databases are required for accurate classification of a significant proportion of PLPs.

Based on our results, we expect first cousins consanguineous couples to be at 16 times higher risk for a child with an AR disorder compared to non-consanguineous couples. This translates to ~3400 newborns with a severe AR disorder per 100,000 births for first cousins. As expected, the risks gradually decreased for more distant relationships and the risk for third cousins was similar to that for non-consanguineous couples at ~0.9%/1.4% for a severe/any AR disorder (**Supplementary Table 6**). These results provide empirical evidence for the common assumption that a third-degree cousin relationship is similar in risk of AR diseases to random mating within an outbred population

For couples in an outbred population, expanding the scope of PCS to wider panels/exome-sequencing is not expected to raise the number of ARCs significantly, due to diminishing marginal returns. A modest number of genes accounts for the majority of ARCs (**Supplementary Fig. 9, Supplementary Table 5**), while genes with rare PLPs hardly impact the ARCs rate. This assessment includes ARCs for variable severity phenotypes. In contrast, consanguineous couples will benefit from a wider scope of PCS due to the significant influence of genes with rare PLPs on the ARCs rate for these couples (**Supplementary Fig. 9, Supplementary Table 5**). Therefore, PCS by extensive gene-panel or exome-sequencing is especially relevant to consanguineous couples.

To assess the effects of consanguinity we devised the CR score, which indicates the increased risk for an AR disorder due to consanguinity. We found that while consanguinity generally increases the risk for an affected child with an AR condition by about 16-fold, this additional risk is not the same for different disorders. Our data show that for consanguineous couples the relative risk for AR-ID and AR skeletal disorders is significantly higher than for other disorders. Whereas about 1 in 3 Dutch individuals carries a PLP variant in a gene for ID, we calculate an expected incidence of only 19 per 100,000 (0.02%) AR ID in offspring of unrelated parents because couples who are both carriers for PLPs in the same gene are rare. For consanguineous couples this rises more than 45-fold to 901 per 100,000 (0.9%). In contrast, for other disorders such as metabolic disorders, the expected incidence is 134 per 100,000 (0.13%) in offspring of unrelated parents, and 1280 per 100,000 (1.3%) in offspring of consanguineous couples. We find that these striking differences are due to differences in the distribution and frequencies of PLPs among different disorders, i.e. differences in their genetic architecture. While in non-consanguineous couples only frequent variants have a strong impact on the ARCs rate, in consanguineous couples even rare variants can have a strong impact on the ARCs rate.

The results in both the Dutch and Estonian cohorts show that ~25% carries a PLP variant in an ID gene, almost all of these variants being rare. There is a single common allele (0.5%) in the Estonian cohort in the *CRADD* gene which is a well-known cause of AR syndromic ID, and likely represents a Northern Scandinavian (Finnish) founder mutation^19^. These observations are in line with previous studies on individuals with ID and other neurodevelopmental disorders (NDD) from outbred populations, which showed a very small (2-3%) contribution of AR variants to ID, with *de novo* pathogenic variants explaining the majority of patients^20,21,22^. In consanguineous couples, a much higher proportion of NDD patients is explained by AR inheritance^20^.

The unique genetic architecture observed for the ID and skeletal disorders compared to other disorders could be explained by a small negative effect on fitness for heterozygous carriers of PLPs in these genes. Our results suggest that there is indeed stronger purifying selection on the ID and skeletal disorders genes with respect to other groups of genes, with selection patterns which are more similar to those of essential genes (**Fig. 5, Supplementary Fig. 8**). Elucidation of the magnitude and mechanisms of such negative fitness effects will require analysis of large population samples with relevant phenotypic read-outs.

We analyzed two distinct Northern European populations, with no geographical relation between them, and found remarkably consistent results. Although these populations have distinct PLPs and common alleles, our estimates of the overall carrier frequency per sample, the most frequently mutated genes, ARCs rates and CRs are very similar. This resemblance may in part be due to shared or similar selection pressures.

This study provides an estimate for the overall burden of AR PLP variants in two European populations. Our approaches can be applied to other populations, in order to establish their specific AR architecture. Such results can be used by clinicians for baseline risk calculations and be incorporated in PCS guidelines. Given that the majority (>85%) of the population carries at least 1 disease allele for any AR disorder, and that 1 in ~4 carries an allele for AR ID, it should now be feasible to study the aggregate effects of these PLPs in terms of development, health, and disease at the population level.

## Methods

### Cohorts

We analysed two European cohorts based on Dutch and Estonian samples. For the purpose of this study, exome data were anonymized. Both populations are from Northern Europe, yet they are distant enough geographically that they can be treated as two distinct European cohorts^11,23^. The Dutch cohort included 4780 unaffected parents of children with intellectual disability (ID), tested by patient-parents trio exome-sequencing. At the time of the analysis, none of the patients was diagnosed with an AR-ID. The exomes were sequenced using DNA isolated from blood, at BGI in Copenhagen, Denmark. Exome capture was performed using Agilent SureSelect v4/v5 and samples were sequenced on an Illumina HiSeq instrument with 101-bp paired-end reads to a median coverage of 75x. Sequence reads were aligned to the GRCh37/hg19 reference genome using BWA version 0.5.9-r16. Variants were subsequently called by the GATK haplotyper (version 3.2-2) and annotated using a custom diagnostic annotation pipeline^24^. The Estonian cohort included 2356 healthy individuals, sequenced as a part of the Estonian Biobank of the Estonian Genome Center, University of Tartu (EGCUT), which is a population-based biobank, containing almost 52,000 samples of the adult population (aged ≥18 years), which closely reflects the age, sex and geographical distribution of the Estonian population. WES samples DNA was enriched for target sequences (Agilent Technologies, Santa Clara, CA, USA; Human All Exon V5+UTRs) according to manufacturer’s recommendations. Sequenced reads were aligned to the GRCh37/hg19 human reference genome using BWA-MEM (version 0.7.7). SAMtools (version 1.2) was applied to compress SAM to BAM (samtools view), sort (samtools sort) and index BAM (samtools index) files. PCR duplicates were then marked using Picard (http://broadinstitute.github.io/picard) (version 1.136) MarkDuplicates.jar. For further BAM improvements, including realignment around known indels and base quality score recalibration, we applied GATK (version 3.4). Single sample genotypes were called by the GATK HaplotypeCaller algorithm (-ERC GVCF).

### Relatedness analysis

We used KING^25^ version 2.2 to calculate kinship coefficients and infer relationships within the cohort. For this analysis, we used 14,643 autosomal variants with a quality score of ≥1000 located in regions covered ≥20x in all samples. KING^25^ inferred 30/52 Dutch/Estonians samples to be related by second degree or closer. Blinded analysis in the Dutch cohort confirmed known relationships in 26 of 30 samples. There was insufficient information to determine relatedness for the other 4 samples, yet they were removed from the analysis since KING^25^ indicated them as having multiple second and third-degree relatives within the cohort.

Overall, we excluded 17 samples from the Dutch cohort and 26 samples from the Estonian cohort by removing one individual from each pair inferred by KING^25^ to have a relationship of second degree or closer.

### Ancestry analysis

a. <u>Dutch population</u> We performed ancestry analysis for the Dutch cohort by LASER^26^ version 2.04, using a reference set of genotyping data from 9,608 loci on chromosome 22 of 700 samples from the Human Genome Diversity Project (HGDP)^27^. The reference samples are subdivided into 7 worldwide populations (Africa, America, Central/South Asia, Europe, Middle East, Oceania, East Asia) and 53 subpopulations. Based on this analysis we excluded 643 samples of non-European descent. Additionally, we analysed the remaining 4120 samples with the ADMIXTURE^28^ tool (version 1.3.0). We used data from the 1000 Genomes Project^18^ to help identify the genetic ancestry of our samples, by using samples from 5 super-populations (Africa, America, Europe, South Asia and East Asia) and running a supervised analysis with K=5. Next, we used the alleles frequencies as an input to a projection analysis for the samples of the cohort. ~97% of the samples had a European component of >0.75, indicating the cohort has homogeneous European ancestry (**Supplementary Fig. 10a**) and confirming the LASER^26^ results.
b. <u>Estonian cohort</u> For the Estonian samples only VCF files were available and therefore we could not run LASER^26^ on this cohort. We ran ancestry analysis by ADMIXTURE^28^, with the same parameters and reference samples as described for the Dutch cohort. Approximately 98% of the samples had a European component of >0.75, indicating the cohort has homogenous European ancestry (**Supplementary Fig. 10b**). Three samples were excluded from the analysis.

After filtering the samples based on ancestry and relatedness, we had 4120 Dutch samples and 2327 Estonian samples of unaffected, European-descent, unrelated individuals that were used for all subsequent analyses in this study.

### Assessing Loss-of-function variants (LoF)

Variants were annotated using the Ensembl Variant Effect Predictor (VEP)^29^ and LOFTEE^30^ tool (Loss-Of-Function Transcript Effect Estimator; Installed at 5th of January 2018, VEP version 91) with the default parameters as a part of the indels filtering process. Indels with Low-confidence score (LC) by LOFTEE^30^ were filtered-out.

### Selection of genes

Genes indicated in OMIM (www.omim.org/) as having both AR and AD phenotypes (AD-AR genes) were assessed by their gnomAD pLI score. This score indicates the probability that a gene is intolerant to a loss of function (LoF) variant. The higher the score the more likely the gene is involved in a dominant disease, and the lower the pLI score, the more likely it is to indicate a recessive disease gene. To determine the appropriate pLI threshold for determining which AD-AR genes are more likely to be AR genes, we generated a reference list of pLI scores for manually curated 930 AR-only genes causing severe phenotypes^7,31^. The 95th percentile score for these genes was 0.86. Therefore, AD-AR genes with a pLI score ≤0.86 were considered as AR and included in subsequent analyses (324 genes).

The final list of AR genes included 1929 genes (6011 transcripts). Of these, OMIM (www.omim.org/) classifies 1605 genes as AR-only phenotypes, and 324 genes as underlying both AR and AD phenotypes. In addition, 1119/1929 genes of the list were manually curated as a part of the development of an Australian PCS panel (Mackenzie’s mission project), and deemed to be associated with severe phenotypes^7^ (**Supplementary Table 1**). Severe was defined in the Australian PCS panel design as: “The condition is one for which an “average” couple would take steps to avoid the birth of a child with that condition”^7^. The Australian PCS project also includes X-linked genes that were not analysed in this study.

### Variant selection and determining variant outcome

We extracted variants located in the exons and flanking 10bp regions of the selected genes, in regions covered ≥20x in ≥90% of the samples. Based on exon positions extracted from Ensembl^29^ (GRCh 37) for the selected genes −94.3% of the coding region±10bp and 58.1% of the exonic regions±10bp (including UTRs) were covered ≥20x in ≥90% of the samples.

We used the list of transcripts from the HGMD database (version 2018.3) (www.hgmd.cf.ac.uk/) as a reference. If a gene had only one transcript described in the HGMD database (www.hgmd.cf.ac.uk/), the outcome of the variant was defined based on this transcript. For genes with several transcripts described in the HGMD (www.hgmd.cf.ac.uk/) database, if >50% of the variant outcomes were LoF/missense/other, this outcome was chosen. All other cases (1777/861 variants in the Dutch/Estonian cohort) were considered as the most severe outcome, and evaluated manually if they passed the selection process.

### Indels filtering

While SNV calling tools generally have good performance, the accuracy of indel calling is relatively low and prone to errors^32^. In order to prevent a high incidence of false positive indels, we adjusted the selection process by using indels in autosomal-dominant genes associated with intellectual disability (AD-ID) (**Supplementary Table 9**) as a proxy for false positives. Since our cohort includes unaffected individuals, indels found in those genes are most probably either not PLPs or false positives. In our Dutch cohort of 4120 samples, there were 1039 indels with ≥500 GATK quality score in 371 AD-ID genes. Raising the quality threshold to ≥1000, removing non-LoF, common (>5% heterozygotes/>1% homozygotes in our cohort; >1% allele-frequency in gnomAD^33^, longer than 10bp, scored LC by LOFTEE^30^, and adjacent indels within 10bp range, decreased the number of AD-ID indels to 46 (4.4%). Of these remaining indels, 12 (26.1%) were in genes with non-LoF pathogenicity mechanism such as known dominant negative/activating PLPs, 20 (43.5%) were in genes with known partial penetrance or variable expression, nine (19.5%) were in genes lacking information about inheritance and pathogenicity mechanism, and five (10.9%) were in genes expected to be affected by LoF variants. Our filtering process thus significantly reduced the number of likely false-positive indels, and was therefore applied to indels in the analysed gene-set.

### Variant selection process

For each cohort, we created a list of presumable PLPs. We included variants with ≥500/1000 GATK quality score for substitutions/indels (respectively), and excluded variants with ≥5% heterozygotes or ≥1% homozygotes frequency within each cohort (**Supplementary Fig. 1**). After manual curation of the frequency drop-outs, we reincluded three known pathogenic variants with >5% carrier frequency in at least one of the cohorts (Dutch/Estonian, respectively): *HFE* p.Cys282Tyr (10.7%/7.6%); *BTD* p.Asp444His (7.1%/8.1%) and *SERPINA1* p.Glu288Val (6.4%/3.2%). This gave rise to 91,341/45,929 variants in total in the Dutch/Estonian cohort, respectively.

We then selected only variants that met at least one of three criteria: (1) Classified as PLP by ClinVar^9^ with a review status of ≥2 stars or classified as PLP by the VKGL^10^ database. This curated database is publicly available and comprises DNA variant classifications established based on (former) diagnostic reports of all 9 Dutch accredited laboratories (www.molgenis.org/vkgl); (2) Loss-of-function (LoF) variants (nonsense, frameshift, canonical splicing) with <1% or unknown frequency in GnomAD^33^. Indels were filtered as described above; (3) Classified as PLP by ≥2/3 databases: InterVar^34^ (an automated ACMG classifier), ClinVar^9^ with a review status of <2 stars, and HGMD (www.hgmd.cf.ac.uk/) (indicated as disease causing variant by the DM flag), and does not contradict the first criterion, i.e. not classified as benign/likely-benign by Clinvar^9^ with a review status of ≥2 stars or by the Dutch database (**Supplementary Fig. 1**).

For the AD-AR genes, only LoF variants were included in the final PLPs list. In total, the selection process filtered out >95% of the initial number of variants in the selected regions (**Fig. 1; Supplementary Fig. 1**).

### Validation of the PLP classification process

Although we used stringent quality scores for the selection of PLPs, we performed several analyses to confirm the validity of our PLP classification process:

1. **Manual classification** Manual classification was performed in 3 groups of PLP variants:

a. <u>PLP Variants with high allele frequency (>1%)</u>. Within the list of 3734/1664 PLPs in the Dutch/Estonian cohort, the majority of variants (3686;98.7%/1613;96.9%), had a carrier frequency of up to 0.05%. There were 16 (0.5%)/18 (1.1%) PLPs with more than 1% carrier frequency in the Dutch/Estonian cohort (**Supplementary Fig. 3**). Amongst these frequent variants, 7 variants were seen with >1% carrier frequency in both cohorts (**Supplementary Table 10**). Manual curation of these variants showed that all common variants were previously described in European populations and/or are known to cause a mild phenotype. The observed frequency was also compared to the frequency reported in the GONL project (http://www.nlgenome.nl/), which is based on sequencing of a different cohort of 498 healthy Dutch individuals (**Supplementary Table 10**). All the variants that were seen at a >1 % frequency in the Dutch cohort were also reported in the GONL database, with 0.2-5.4% allele frequency (mean 1.6%).
b. <u>PLP variants in a homozygous state</u> For both the Dutch/Estonian cohorts, only 42/11 (1%/0.5%) of the samples were homozygous for any of the PLPs. This percentage was even lower for the set of severe recessive genes (12/4; 0.3%/0.2%). Overall, 19 PLPs were seen in a homozygous state in one sample or more. Manual curation of these variants showed that 11 of these have been reported to cause only a mild phenotype or even appear asymptomatic when seen in a homozygous state, and 6 have conflicting evidence about their pathogenicity.
c. <u>Curation of the 214 PLPs in genes underlying deafness</u> All PLPs found in deafness genes were manually classified by an expert who used, among other databases, the Deafness Variation Database (DVD), a comprehensive, open-access resource that integrates all available genetic and clinical data together with expert curation^35^. Of the 214 variants our selection process classified as PLP, expert manual curation classified one variant as likely-benign, 6 variants were as VUS, 174 variants as LP and 33 variants as P. Overall, 96.7% (207/214) of the variants were correctly classified as PLPs.
2. **Assessing selection process performance for PLP variants in *CFTR*** We extracted the list of variants that are classified as PLP in the CFTR2 database (cftr2.org; version Jan10-2020) and ran it through the selection process. The selection process classified correctly as PLP 347/414 (83.8%) variants, including 113 missense variants. Most of the variants that were not classified as PLP (59/67; 88%) were missense variants. Six variants were non-coding non-canonical splice site region variants, one variant was an inframe insertion, and one variant was an intronic variant.
3. **Analysis of selected missense variants** In order to assess the pathogenicity of missense variants classified as PLP based on tier 3 criteria of the selection process, we compared CADD scores of missense variants from tier 1 and tier 3 with the CADD scores of those which failed to pass the selection process (non-PLP). The CADD scores of missense variants classified as PLPs based on tier 3 criteria are similar to those of missense variants classified as PLPs based on tier 1 criteria, and are significantly higher than those of missense non-PLPs (**Supplementary Fig. 11**).

### Virtual matings

We simulated all possible matings within the 4120 samples of the Dutch cohort (8,485,140 theoretical couples), and the 2327 samples of the Estonian cohort (2,706,301 theoretical couples), irrespective of gender. Variants that were manually classified for high frequency/homozygosity, and were proven to be asymptomatic or cause a very mild phenotype in the homozygous state, were excluded if the virtual mating was predicted to be at-risk for a homozygous offspring, yet included if the virtual mating was predicted to be at-risk for a compound heterozygous offspring (**Supplementary Table 4**).

The ARCs rates were computed by simulations rather than using allele frequencies. This method is the most accurate way to assess ARCs rates since it is based on actual genotypes from the population, whereas the calculation using allele frequencies necessarily needs to assume linkage equilibrium between all variants.

### Pathogenic variants simulations

The number of shared PLPs between the Dutch and Estonian cohorts was 373 (10% of the Dutch PLPs and 22% of the Estonians PLPs) (**Fig. 2a**). In order to determine whether this proportion of shared variants is significantly less than expected, we ran 10000 simulations in which we randomly created two groups of PLPs chosen from the list of 3734 Dutch PLPs. The sizes of the groups were 2582 and 1152, keeping the ratio of 2.24 that was seen for the Dutch vs Estonian PLPs (3734/1664). For each simulation we checked the proportion of shared PLPs between the two random groups. The mean number of shared variants was 799 (21.4%).

### Gene rankings

Genes in which PLPs were observed were ranked by the frequency of PLPs observed. Six genes were included in the top-10 rankings of PLPs per gene in both cohorts (**Fig. 2b**). In order to check whether this number of genes in the top-10 rankings is statistically significant, we used Fisher exact test (P-value= 1·10^-5^).

### Consanguinity analysis

For the consanguinity calculations, we verified that variant frequencies in the two populations adhere to the Hardy-Weinberg equilibrium. We compared the observed number of WT (AA), heterozygotes (Aa), and homozygotes (aa) to the expected numbers (pp,2pq,qq) using a chi-squared test, showing that variants in both populations adhere to the Hardy-Weinberg equilibrium and there is no significant difference between observed and expected with a P-value of 0.4 for the Dutch cohort and 0.1 for the Estonian cohort. Next, we calculated the expected risk for different degrees of consanguinity, relying on the expected proportion of shared alleles in each relationship: 1/8 for first cousins, 1/16 for first cousins once removed, 1/32 for second cousins, and 1/128 for third cousins. Therefore, the probability of a couple to be at risk in a given gene was calculated as 2pq*<expected proportion of shared alleles>*cohort size, where q is the sum of allele frequencies of PLP variants in a given gene, and p=1-q.

### Gene panels

All 1929 genes were divided into panels based on their related disorders (**Supplementary Table 1**). The ID-related genes were divided into two panels-ID and Metabolic-ID. The ID panel includes genes that are related to syndromic and non-syndromic ID, but does not include metabolic genes. All genes that are related to metabolic disorders with or without phenotypes other than ID were included in the metabolic panel. An additional gene group, multisystem disorders, comprised all genes underlying more than one phenotype, excluding intellectual-disability (ID) and metabolic disorders (**Supplementary Table 1**).

### Consanguinity ratio (CR)

In order to examine whether the differences between the CR scores are statistically significant, we ran 5000 simulations per panel for each cohort. In each simulation, we randomly assigned genes into panels of same total bp length as the original panels (**Supplementary Fig. 6**). We computed the P-values based on how many times the simulated CR was different from the actual CR for each panel using a twotailed test with a Bonferroni correction (**Supplementary Fig. 6; Supplementary Table 12**).

### Analysis for genetic selection

To determine whether groups of genes show different patterns of selection we used three different scores that serve as a proxy of purifying selection. We calculated the normalized gene singleton density, the coding regions singleton density (coding singleton density)^36^, and the Residual Variation Intolerance Score (RVIS)^37^ for Europeans samples on the 1000 genomes data^18^ (**Supplementary Table 13**).

We compared the scores of gene sets described in this study to reference lists of genes that show different signatures of purifying selection: “Essential” genes^38^ and “haploinsufficiency severe” genes^39^ as target of strong purifying selection, and genes labeled as “Non Essential”^38^ and “Olfactory” receptors^40^ as controls (no strong selection signatures) (**Supplementary Table 13**). Statistical significance of the differences between the medians was evaluated using the Wilcoxon-rank sum test for the gene panel ID/Skeletal compared to all other disorders (**Fig. 5, Supplementary Fig. 8**).

Next, we simulated a large meta-population with ten subpopulations with an effective population size (Ne) of 10000 each. Each population could exchange migrants with a rate of 1% in each generation. We simulated the possibility of a locus to mutate and its increase in frequency in different scenarios: no selection (s=0) and increased purifying selection against the heterozygotes (s>0), with a total negative selection against homozygotes (s=1). The simulated selection coefficients are linked to the percentage of reduction of an individual to have offspring (for s=0.05 heterozygous carriers have 5% less probability to mate and have offspring). The simulations were done using simuPOP^41^ version 1.1.7 (**Supplementary Fig. 7**).

## Acknowledgements

C.T.S. and Y.X. were supported by Wellcome grant number 098051. A.M. was partially supported by the EU through the ERD Fund, Project No. 2014-2020.4.01.15-0012) “Gentransmed”. S.C. thanks the Israel Science Foundation grant 407/17 and the Abisch-Frenkel Foundation.

Inclusion of RadboudUMC data was in part supported by the Solve-RD project that has received funding from the European Union’s Horizon 2020 research and innovation program under grant agreement no. 779257. This work was in part financially supported by grants from the Netherlands Organization for Scientific Research (917-17-353 to C.G.).

We thank Maartje van de Vorst and Karolis Sablauskas for helping with data analysis.

## Author Contributions

H.F., E.L.L., C.G., and H.G.B. contributed to the generation of figures and writing of the manuscript.

R.M., R.A., A.M, and H.G.Y. contributed to the generation and quality control of data. M.M. developed methods, contributed data or performed analyses.

H.G.Y., C.T. Y.X., S.C., E.L.L., C.G. and H.G.B. provided experimental and analytical supervision.

E.L.L., C.G. and H.G.B. provided project supervision.

## Data availability statement

All variants that were classified as PLP in this study are listed in Supplementary Table 14.

## Code availability statement

Scripts and code are available upon request.

## Ethical regulations

Dutch exomes were obtained as a part of an anonymized study, following guidelines for anonymized study procedures, based on approval by the institutional review board ‘Commissie Mensgebonden Onderzoek Regio Arnhem-Nijmegen’ under number 2011/188.

Estonian exomes were obtained as a part of the Estonian Biobank project. permission nr 234/T - 12 to study human exomes in the Estonian Biobank. All subjects in the Estonian Biobank have signed informed consent and the whole biobanking program is governed by special law: “Human Genes Research Act”.

