## Supplemental figures and tables for "The landscape of autosomal-recessive pathogenic variants in European populations reveals phenotype-specific effects"

**Supplementary Figures**

**Supplementary Figure 1**. **The variant classification process and number of PLPs identified in each cohort**

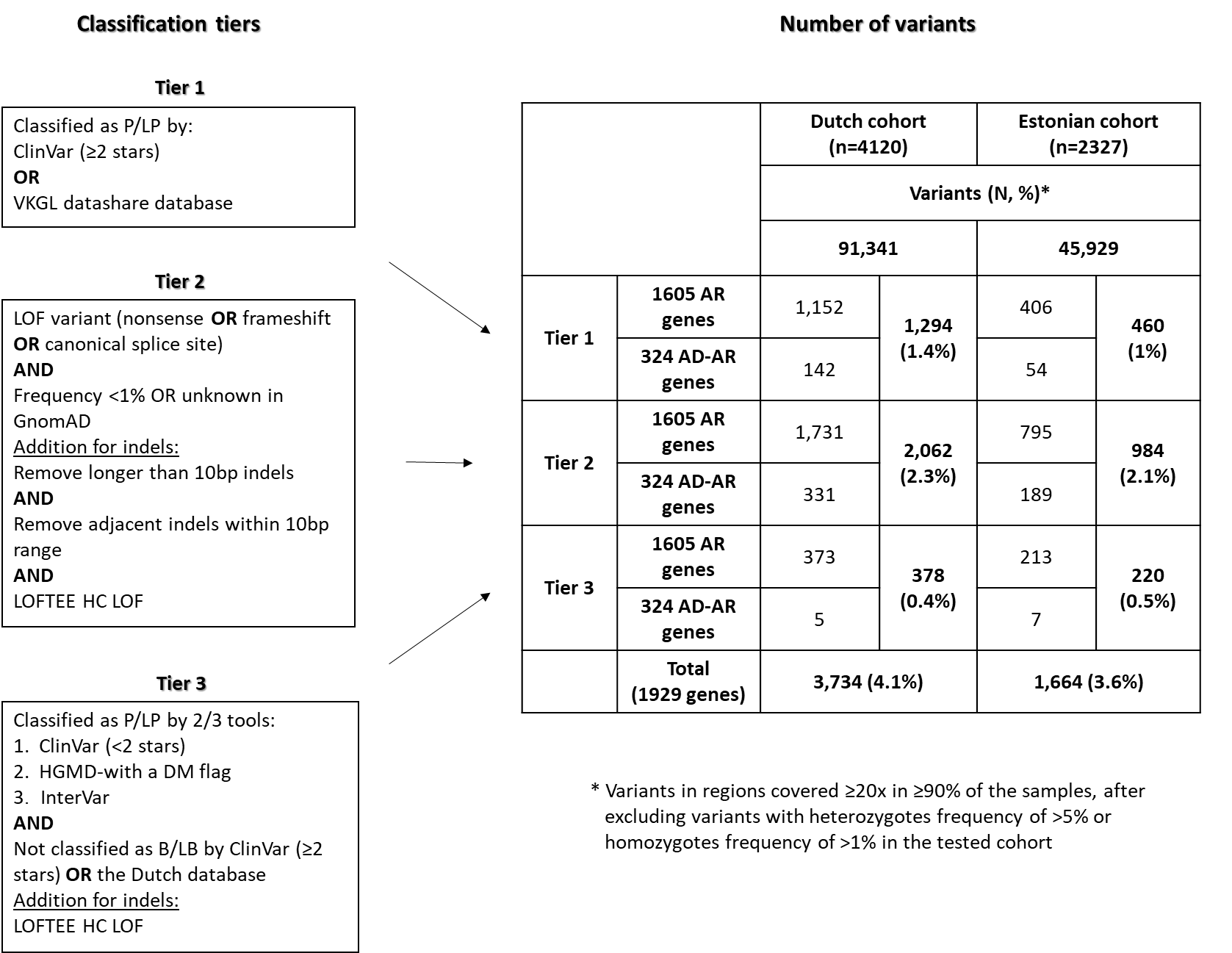

**Supplementary Figure 2**. **Distribution of the number of heterozygous PLP variants per sample**
The carrier distribution is presented for the Dutch **(a)** and Estonian **(b)** cohorts for 1929 AR genes
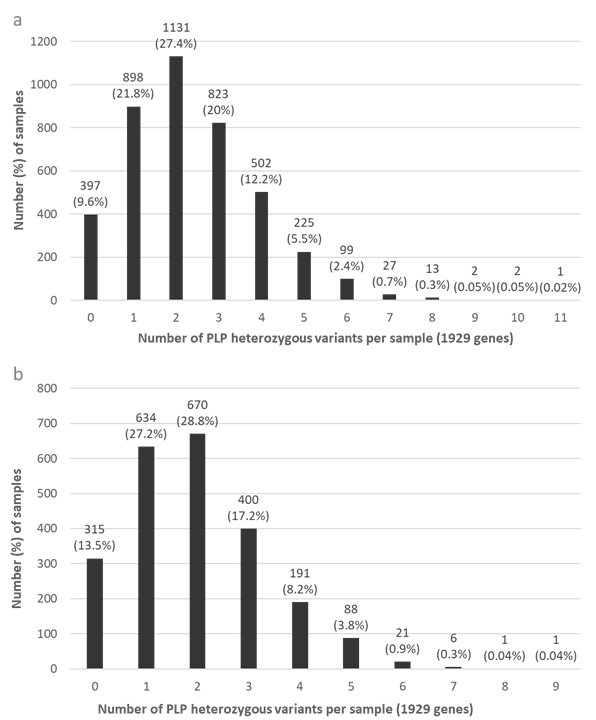

**Supplementary Figure 3**. **Distribution of heterozygous PLP variants by carrier frequency**
The PLPs distribution is presented for the Dutch **(a)** and Estonian **(b)** cohorts for 1929 AR genes

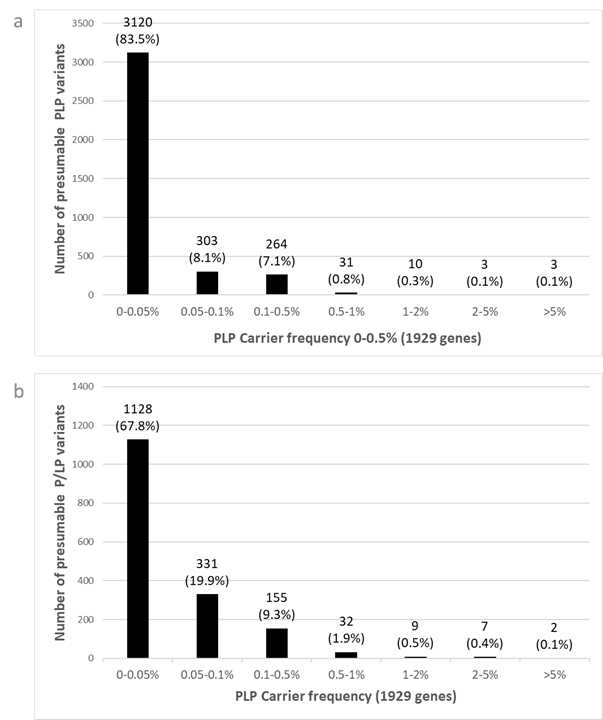

**Supplementary Figure 4**. **Distribution of heterozygous PLP variants per gene** **by carrier frequency**
The distribution of PLPs per gene is presented for the Dutch **(a)** and Estonian **(b)** cohorts for 1929 AR genes.

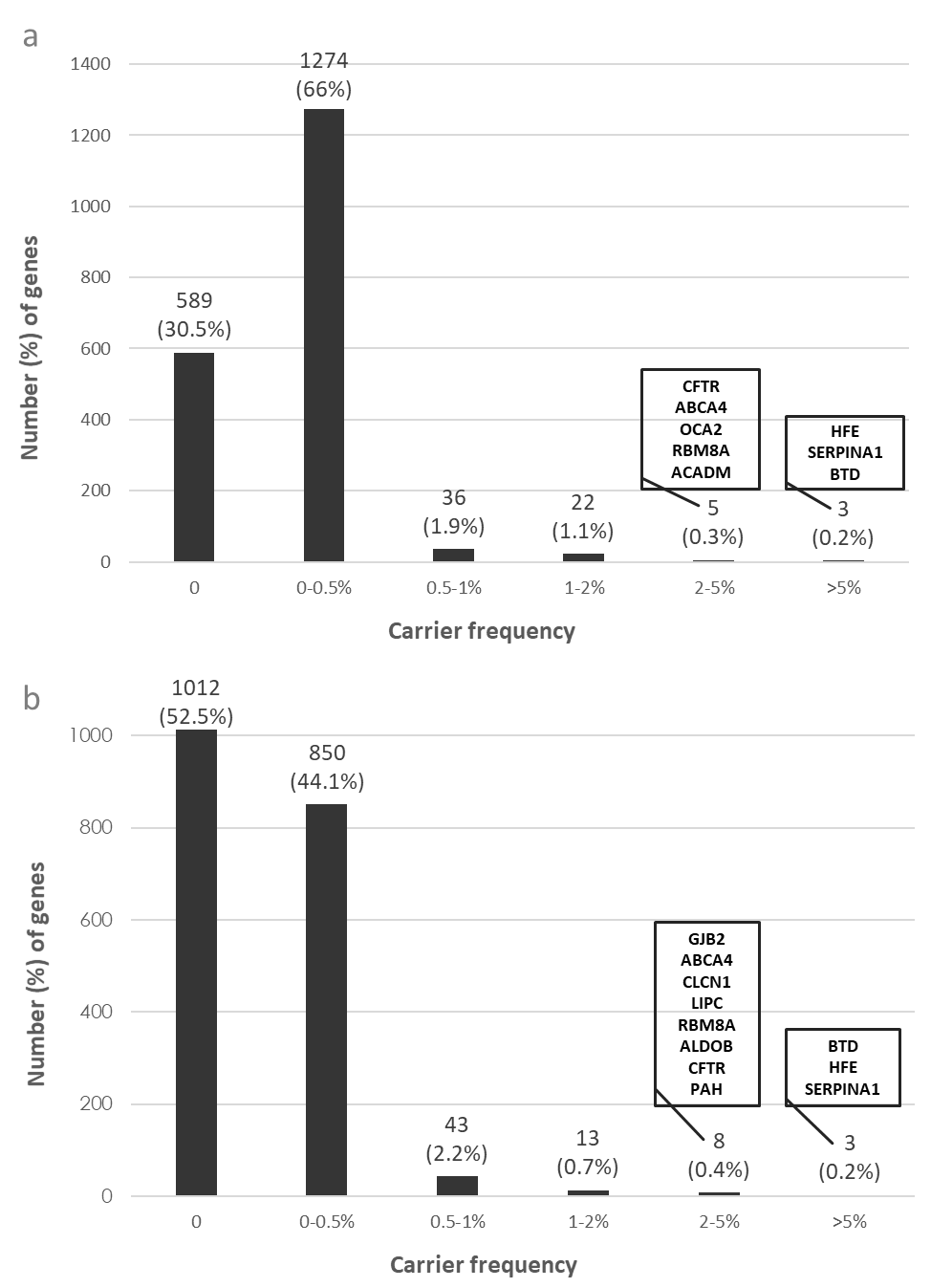

**Supplementary Figure 5**. **Distribution of genes by allele-counts in different disorders**

Distribution of the total allele-counts (number of heterozygotes added to twice the number of homozygotes) in the **(a)** Dutch cohort and **(b)** Estonian cohort. Different colors represent different allele-count ranges.

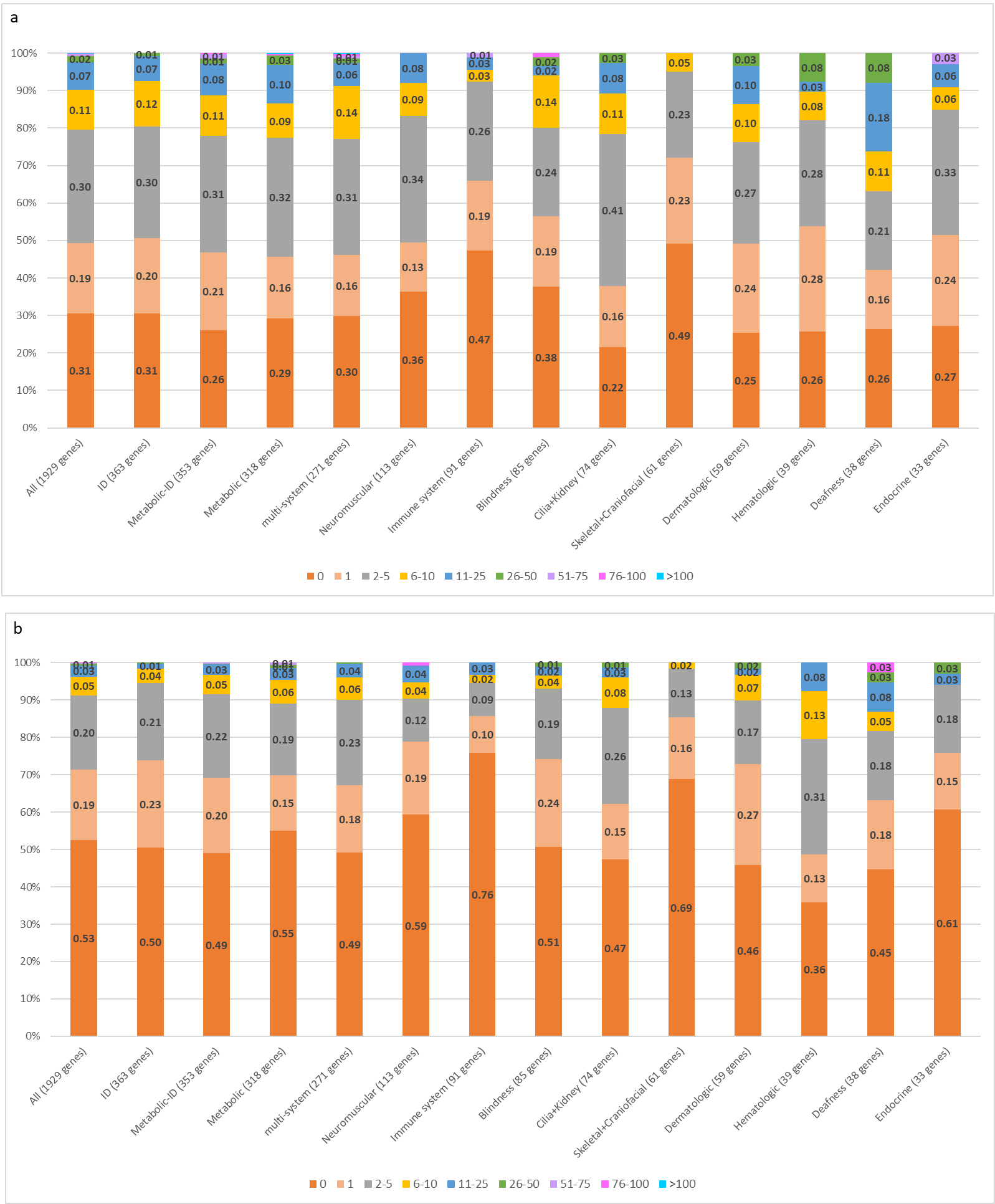

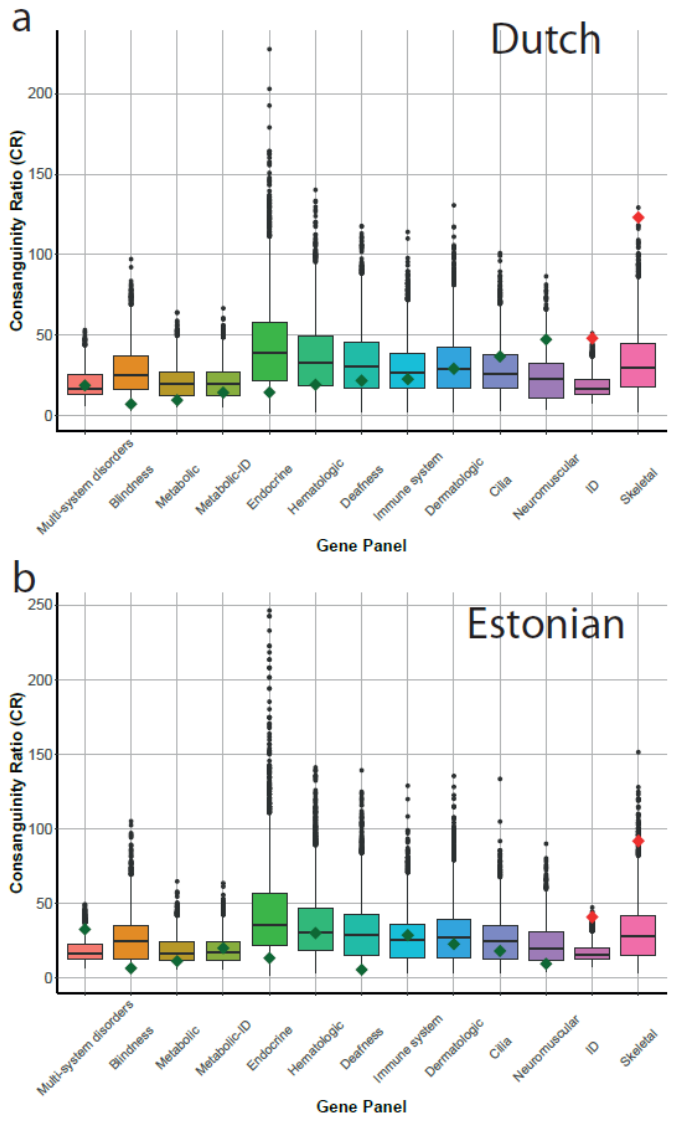
**Supplementary Figure 6**. **Permutations of consanguinity ratio (CR) based on coding length of the gene panels**
Results are shown for the **(a)** Dutch cohort and **(b)** Estonian cohort. Boxplots indicate the distribution of simulated values for each gene panel. Actual values are indicated by diamonds where colors are as follows: Green: no significant difference; Red: actual CR is significantly higher. P-values were calculated for a two-tailed test. Nominal P-values are 4·10^-3^ in both cohorts for the ID panel, and 3·10^-3^/0.02 in the Dutch/Estonian cohort for the skeletal panel (**Supplementary Table 12**).

**Supplementary Figure 7. Simulations of the impact of selection effects on the frequency of genetic variants in the population**

We simulated a large population system based on ten subpopulations with an effective population size (Ne) of 10,000 each (**Methods**). The points represent the average allele frequency across 100 replicates for each scenario. Absence of selection lead to an increase of the mutation frequency up to 0.2% in 1000 generations, a relatively weak selection pressure of 0.001 against heterozygous individuals reduce the mutation frequency to 0.17%, to 0.09% with s=0.005, to 0.05% with s=0.01 and to 0.01% with a stronger negative selection of s=0.05. In particular we observed no significant change of frequency after 200 generations for scenarios with selection coefficients >0.005.

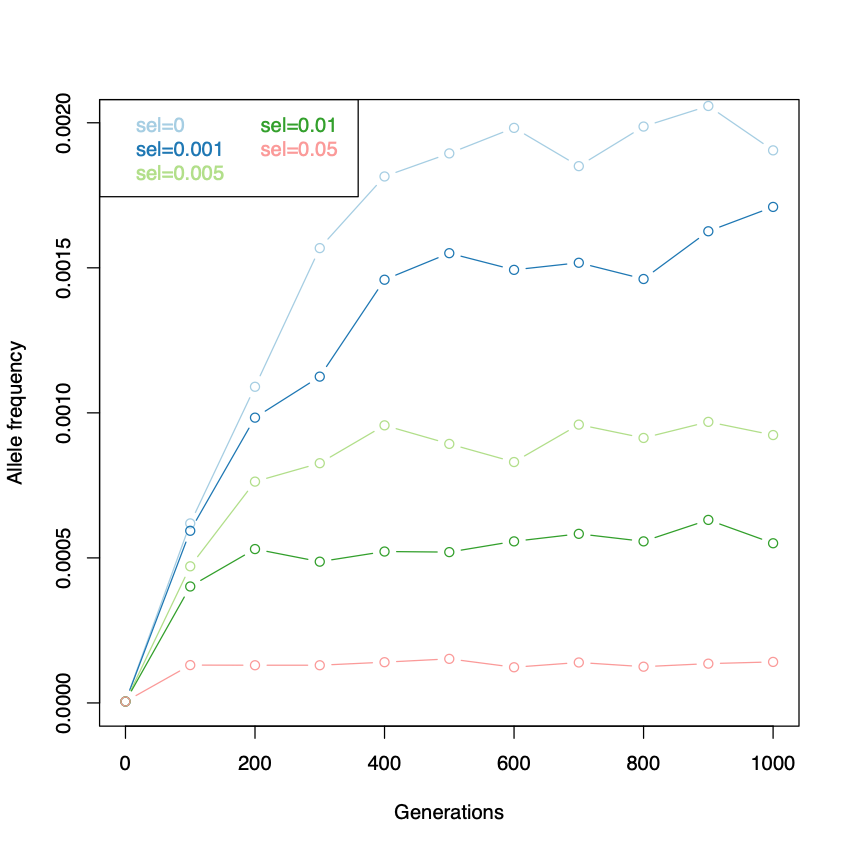

**Supplementary Figure 8. Metrics of natural selection for the different gene panel categories based on 1000 genomes dataset**

Violin plots of the distribution of gene singleton density scores **(a)** and RVIS **(b)** for different gene panels (**Supplementary Table 13**). Gray colors indicate external reference gene sets, whereas colored violin plots are gene sets as defined in this manuscript. Tested differences of the median are indicated by braces.

**(a) Gene singleton density**. Mean gene singleton density was significantly different for ID/skeletal compared to other disorders, using a Wilcoxon rank sum test (P-value- 6.5·10^-5^).

**(b) RVIS for the different panels.** Mean RVIS was significantly different for ID/skeletal compared to other disorders, using a Wilcoxon rank sum test (P-value- 3.6·10^-6^).

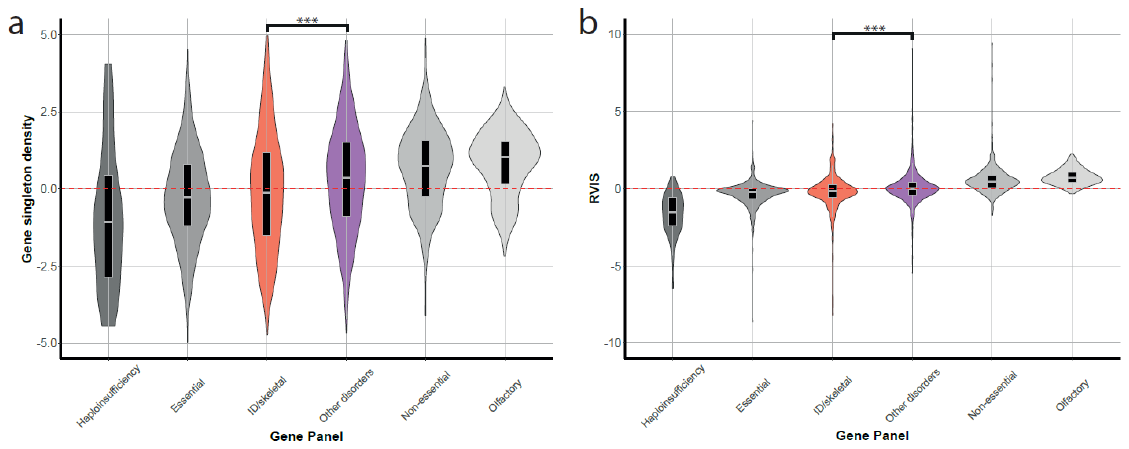

**Supplementary Figure 9. Relationship between the number of genes needed to explain a fraction of ARCs for different gene panels**

The X-axis is the number of genes that explain a fraction of ARCs, ordered by decreasing carrier frequency. The Y-axis is the fraction of ARCs due to PLP variants in these genes. Red lines show the number of genes in a non-consanguineous population, and green lines for first cousins. Dotted lines indicate the 90% of ARCs explained for non-consanguineous individuals and first cousins. **(a)** Dutch cohort. **(b)** Estonian cohort. Numbers for all genes and severe genes are presented at **Supplementary Table 5.**

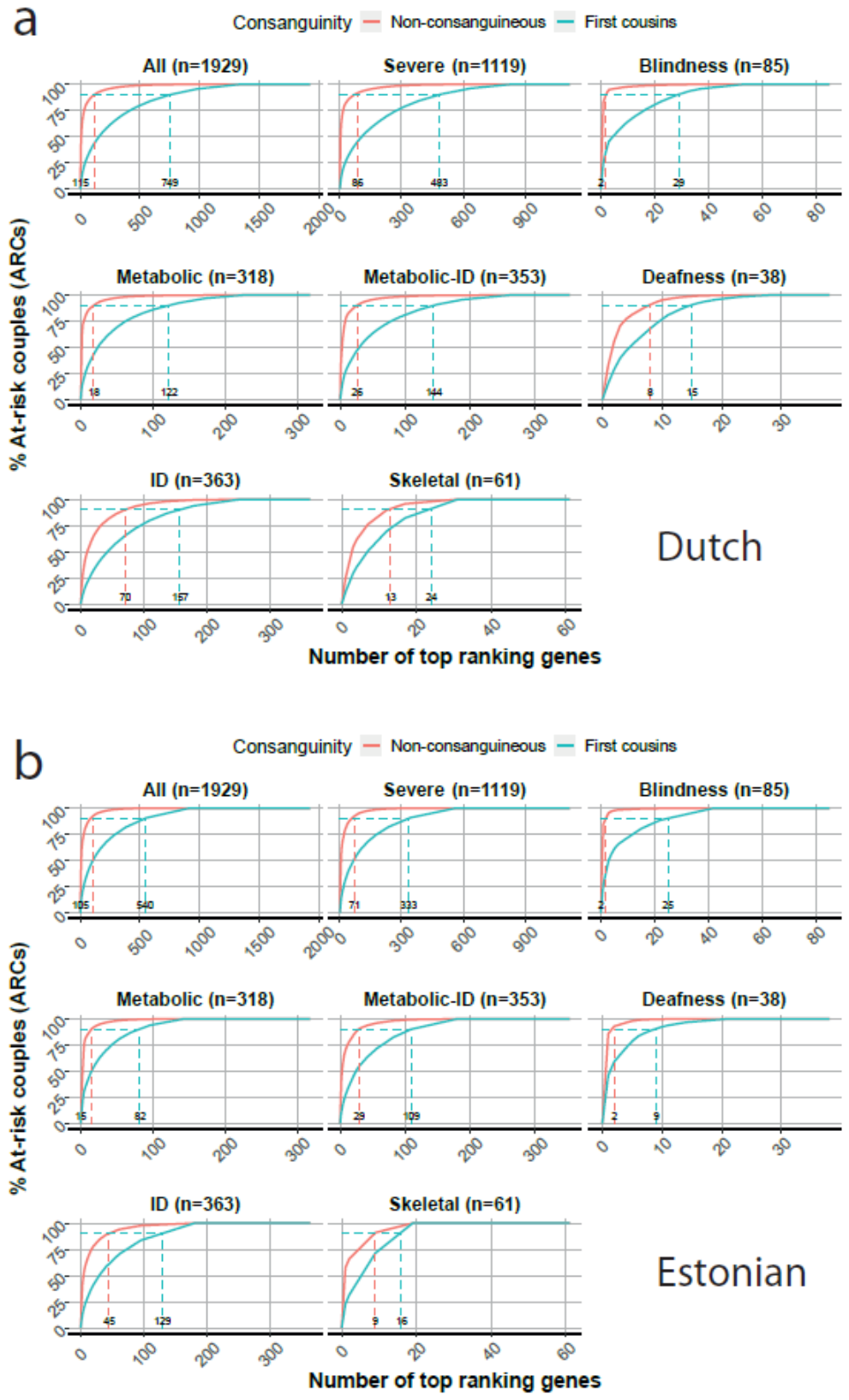

**Supplementary Figure 10**. **Ancestry analysis of the Dutch and Estonian cohort**
The analysis was done by ADMIXTURE projection analysis for the **(a)** Dutch and **(b)** Estonian cohort using K=5

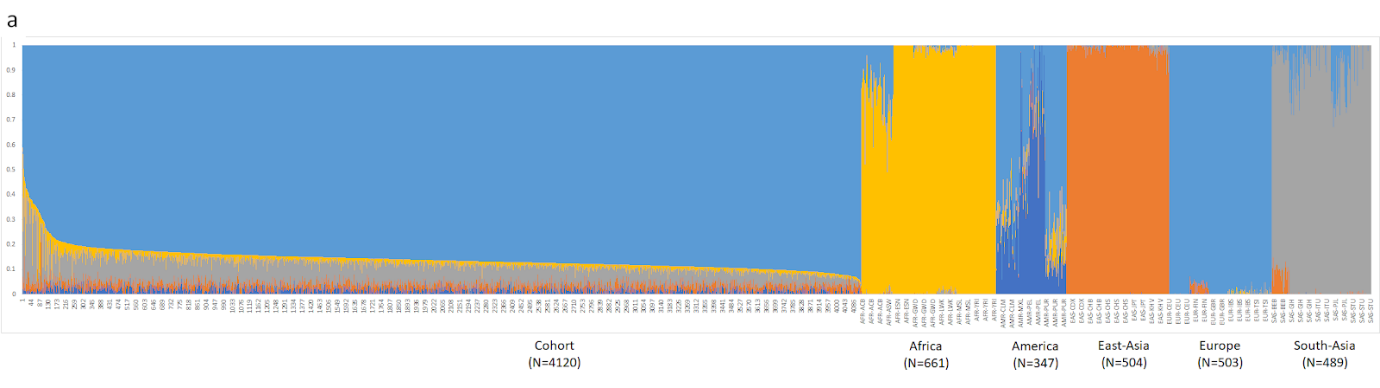

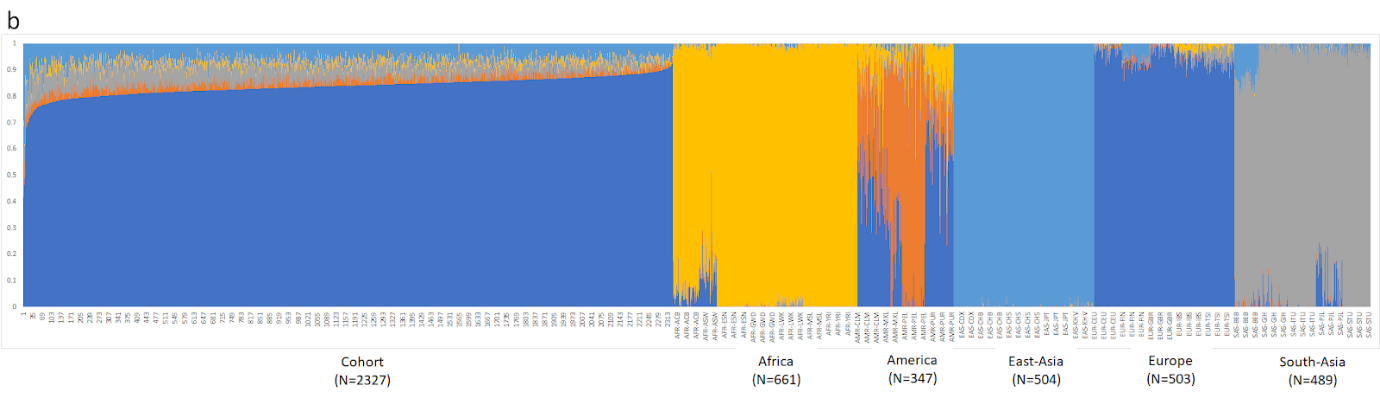

**Supplementary Figure 11**. **Validation of tier 3 classification criteria: CADD score comparison**

Box plot of the CADD scores of variants classified as PLP using tier 1 or tier 3 criteria and variants classified as non-PLP (**Supplementary Fig. 1**); The CADD scores of variants classified as PLP using either tier 1 or tier 3 criteria were similar (P=0.97), whereas the CADD scores of either tier 1 or tier 3 PLPs were significantly higher than those of non-PLP (P=2.2·10^16^ and P=2.2·10^16^ respectively), using Wilcoxon signed rank test.

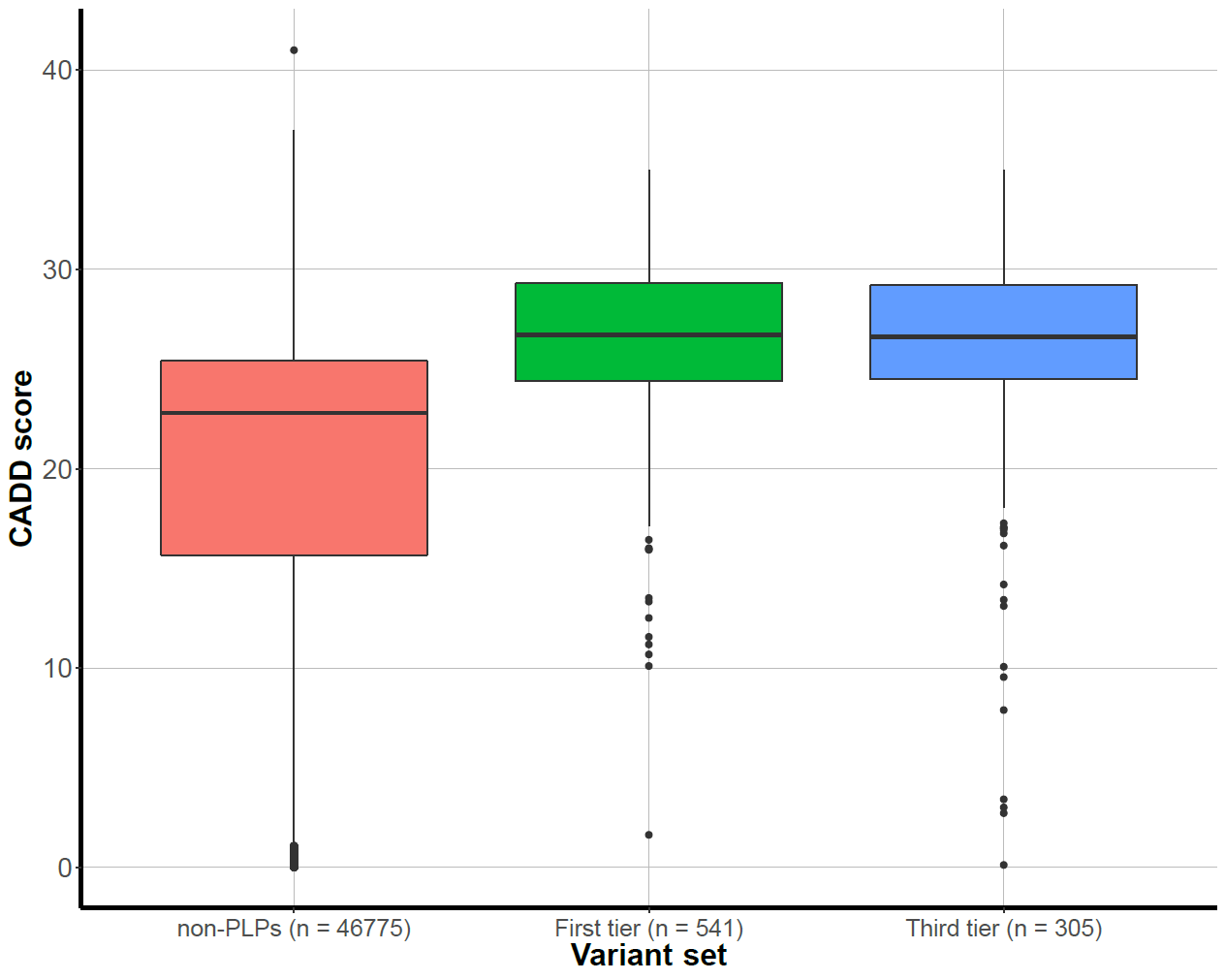

**Supplementary Tables**

**Supplementary Table 1**. **List of the 1929 analysed genes (excel)**

The table presents gene names, transcripts that were analysed, OMIM ID, rather the gene is defined as related to a severe phenotype and which gene-panel it belongs to.

**Supplementary Table 2**. **Genes with ≥1% carrier frequency in the Dutch and the Estonian cohorts**

Dark shadow= genes with ≥1% carrier frequency in both cohorts

Light shadow= genes with ≥1% carrier frequency in one cohort and ≥0.5% in the other cohort

| **Gene** | **Severe phenotype** | **Dutch cohort** | | **Estonian cohort** | |
| --- | --- | --- | --- | --- | --- |
|  |  | **Number of heterozygotes**  **(N/4120)** | **Number of homozygotes**  **(N/4120)** | **Number of heterozygotes (N/2327)** | **Number of homozygotes**  **(N/2327)** |
| *HFE* | - | 442 | 18 | 176 | 3 |
| *SERPINA1* | V | 434 | 8 | 159 | 2 |
| *BTD* | - | 327 | 3 | 193 | 2 |
| *CFTR* | V | 187 | 0 | 53 | 0 |
| *ABCA4* | V | 171 | 0 | 90 | 2 |
| *OCA2* | V | 101 | 0 | 4 | 0 |
| *RBM8A* | - | 93 | 3 | 63 | 0 |
| *ACADM* | - | 90 | 0 | 5 | 0 |
| *MUTYH* | - | 83 | 0 | 19 | 0 |
| *PAH* | V | 79 | 0 | 51 | 0 |
| *PMM2* | V | 75 | 0 | 15 | 0 |
| *ACY1* | - | 71 | 0 | 25 | 0 |
| *USH2A* | V | 69 | 0 | 20 | 0 |
| *GALC* | V | 67 | 0 | 10 | 0 |
| *GNRHR* | - | 67 | 1 | 35 | 0 |
| *MPO* | - | 63 | 0 | 12 | 0 |
| *CBS* | V | 59 | 0 | 8 | 0 |
| *DYNC2H1* | V | 56 | 0 | 27 | 0 |
| *DHCR7* | V | 55 | 0 | 15 | 0 |
| *TMPRSS3* | - | 48 | 0 | 10 | 0 |
| *ABCG8* | - | 47 | 0 | 12 | 0 |
| *HEXA* | V | 47 | 0 | 3 | 0 |
| *TF* | V | 47 | 1 | 19 | 0 |
| *NAGA* | V | 45 | 0 | 10 | 0 |
| *SI* | - | 44 | 0 | 15 | 0 |
| *ATP7B* | V | 44 | 1 | 10 | 0 |
| *TGM5* | - | 43 | 0 | 32 | 0 |
| *CNGB3* | V | 43 | 0 | 21 | 0 |
| *FGB* | V | 43 | 0 | 4 | 0 |
| *ALDOB* | V | 42 | 0 | 56 | 0 |
| *AGXT* | V | 40 | 0 | 26 | 0 |
| *GJB2* | - | 32 | 0 | 97 | 1 |
| *NBN* | V | 30 | 0 | 26 | 0 |
| *LIPC* | V | 25 | 0 | 71 | 0 |
| *CLCN1* | - | 78 | 0 | 20 | 0 |
| *UGT1A1* | V | 43 | 0 | 10 | 0 |
| *UPB1* | V | 19 | 0 | 28 | 0 |
| *EDN3* | - | 14 | 0 | 32 | 0 |
| *CYP24A1* | - | 14 | 0 | 24 | 0 |
| *OTOG* | - | 12 | 0 | 28 | 0 |
| *PKD1L1* | - | 12 | 0 | 24 | 0 |
| *NUP93* | V | 7 | 0 | 41 | 0 |

**Supplementary Table 3**. **Neonatal screening and data-based estimations comparison**

Children that are predicted to be homozygous for a mild variant in the *CFTR*/*BTD* genes were excluded (**Supplementary Table 4**), yet compound heterozygotes with another PLP variant were not. Spearman correlation coefficient for both years: 0.99
* Excluding *CFTR* p.R117H and *BTD* p.D444H

| **Gene/s** | **Disease** | **Our data** | | **Screening 2016** | | **Screening 2017** | |
| --- | --- | --- | --- | --- | --- | --- | --- |
|  |  | **Expected* number of affected children for N=170,000** | **Rank*** | **Observed number of affected children (N=172,754)** | **Rank** | **Observed number of affected children (N=169,883)** | **Rank** |
| *ACADM* | Medium-chain acylCoA dehydrogenase deficiency | 20.1 | **3 (2)** | 1 | **5** | 18 | **2** |
| *ACADVL* | Very long chain acyl-CoA dehydrogenase deficiency | 1.4 | **6 (6)** | 3 | **3** | 4 | **5** |
| *BCKDHA/BCKDHB/DBT* | Maple syrup urine disease | 0.1 | **11 (11)** | 1 | **5** | 0 | **9** |
| *BTD* | Biotinidase deficiency | 50.7 (2.6) | **2 (4)** | 1 | **5** | 9 | **4** |
| *CFTR* | Cystic-fibrosis | 80.9 (50.8) | **1 (1)** | 29 | **1** | 23 | **1** |
| *FAH* | Tyrosinaemia type 1 | 0.3 | **10 (10)** | 1 | **5** | 2 | **7** |
| *GALT* | Galactosemia | 2.3 | **5 (5)** | 0 | **6** | 0 | **9** |
| *GCDH* | Glutaric aciduria type 1 | 0.5 | **9 (9)** | 1 | **5** | 0 | **9** |
| *HADHA/HADHB* | Trifunctional Protein deficiency / Long-chain hydroxyacyl-CoA dehydrogenase deficiency | 0.5 | **9 (9)** | 0 | **6** | 1 | **8** |
| *IVD* | Isovaleric aciduria | 0.8 | **7 (7)** | 2 | **4** | 3 | **6** |
| *MCCC1/MCCC2* | 3-Methylcrotonyl-CoA carboxylase deficiency | 0.6 | **8 (8)** | 2 | **4** | 3 | **6** |
| *PAH* | Phenylketonuria | 15.4 | **4 (3)** | 16 | **2** | 12 | **3** |

**Supplementary Table 4**. **Variants that were excluded from the ARCs calculations for an expected child in a homozygous state**

| **Gene** | **Chromosome** | **Position (GRCh37/hg19)** | **Reference** | **Alternation** |
| --- | --- | --- | --- | --- |
| *ABCA4* | chr1 | 94473807 | C | T |
| *ABCA4* | chr1 | 94517254 | C | G |
| *RBM8A* | chr1 | 145507646 | G | A |
| *BTD* | chr3 | 15686693 | G | C |
| *FGB* | chr4 | 155489608 | C | T |
| *HFE* | chr6 | 26093141 | G | A |
| *CFTR* | chr7 | 117171029 | G | A |
| *GALC* | chr14 | 88452941 | T | C |
| *SERPINA1* | chr14 | 94844947 | C | T |
| *SERPINA1* | chr14 | 94847262 | T | A |
| *STRC* | chr15 | 43892808 | T | G |
| *EDN3* | chr20 | 57897443 | G | GA |

**Supplementary Table 5. Number of genes that accounts for ARCs**

The numbers represent N most mutated genes that accounts for %ARCs

| **%ARC** | **1929 genes** | | | | **1119 severe genes** | | | |
| --- | --- | --- | --- | --- | --- | --- | --- | --- |
|  | **Non-consanguineous- explained by N most frequent genes** | | **Consanguineous- explained by N most frequent genes** | | **Non-consanguineous- explained by N most frequent genes** | | **Consanguineous- explained by N most frequent genes** | |
|  | **Dutch** | **Estonian** | **Dutch** | **Estonian** | **Dutch** | **Estonian** | **Dutch** | **Estonian** |
| 10% | 1 | 1 | 10 | 6 | 1 | 1 | 6 | 5 |
| 20% | 2 | 2 | 27 | 19 | 1 | 2 | 17 | 13 |
| 30% | 3 | 3 | 56 | 38 | 2 | 3 | 37 | 25 |
| 40% | 5 | 5 | 98 | 66 | 3 | 4 | 66 | 42 |
| 50% | 8 | 8 | 156 | 105 | 4 | 6 | 107 | 67 |
| 60% | 13 | 11 | 235 | 160 | 6 | 8 | 161 | 99 |
| 70% | 24 | 21 | 344 | 241 | 12 | 15 | 231 | 148 |
| 80% | 46 | 39 | 504 | 358 | 23 | 27 | 330 | 220 |
| 90% | 115 | 84 | 749 | 540 | 70 | 57 | 483 | 333 |
| 100% | 980 | 552 | 1339 | 915 | 637 | 341 | 833 | 564 |

**Supplementary Table 6**. **ARCs rates for different levels of consanguinity and different gene-sets in both the Dutch and Estonian cohorts**

|  | **1929 genes** | | **1119 severe genes** | |
| --- | --- | --- | --- | --- |
|  | **Dutch** | **Estonian** | **Dutch** | **Estonian** |
| **1st cousins** | 24.90% | 20.91% | 16.42% | 13.37% |
| **1st cousins once removed** | 12.45% | 10.45% | 8.21% | 6.69% |
| **2nd cousins** | 6.22% | 5.23% | 4.11% | 3.34% |
| **3rd cousins** | 1.56% | 1.31% | 1.03% | 0.84% |
| **Non- consanguineous** | 1.47% | 1.28% | 0.99% | 0.76% |

**Supplementary Table 7**. **Comparison to manually classified PLP variants (Dutch National Clinical Genetics laboratories (VKGL) database)**
ID= intellectual disability

|  | **Deafness** | **Blindness** | **ID** | **Metabolic** |
| --- | --- | --- | --- | --- |
| **Number of PLPs in the Dutch VKGL database in 1605 genes with  AR-only phenotype** | 364 | 968 | 1752 | 1592 |
| **+ SNV** | 310/364  (85.2%) | 722/968 (74.6%) | 1368/1752 (78.1%) | 1328/1592 (83.4%) |
| **+ In regions of 20x coverage in >=90% of the samples** | 280/310 (90.3%) | 630/722 (87.2%) | 1206/1368 (88.1%) | 1182/1328 (89%) |
| **+ Will be classified as PLP** | 220/280  (**78.6%**) | 518/630 (**82.2%**) | 924/1206 (**76.6%**) | 816/1182 (**69%**) |

**Supplementary Table 8**. **Cumulative number of PLPs per sample and ARCs for a severe phenotype for each tier of the analysis**

| **Filtering step** | | **Dutch cohort (N=4120)** | | **Estonian cohort (N=2327)** | |
| --- | --- | --- | --- | --- | --- |
|  |  | **# of PLPs per sample** | **ARCs rate-  1119 genes (x/8,485,140)** | **# of PLPs per sample** | **ARCs rate-**  **1119 genes  (x/2,706,301)** |
| **Tier 1** | **ClinVar with >=2 stars** | 0.8 | 58,042 (0.7%) | 0.8 | 9,550 (0.8%) |
|  | **+ Dutch VKGL database** | 1.3 | 89,560 (1%) | 1 | 24,038 (0.9%) |
| **Tier 2** | **+Rare LOF** | 2 | 101,714 (1.2%) | 1.6 | 27,231 (1%) |
| **Tier 3** | **+ >=2/3 tools** | 2.3 | 124,722 (1.5%) | 2 | 34,570 (1.3%) |

**Supplementary Table 9**. **Filtering process for indels**
The filtering of indels is based on indels analysis in 371 intellectual disability autosomal-dominant genes (**Methods**).

| **Filtering step** | **Remaining indels (N(%))** |
| --- | --- |
| Quality score of ≥500 and coverage ≥20x for all samples | 1039 |
| Quality score of ≥1000 and coverage ≥20x for all samples | 599 (57.6%) |
| LoF variants (frameshifts, stop-gained, stop-lose) | 97 (9.3%) |
| Excluding common variants (>5% heterozygotes/>1% homozygotes in our cohort; >1% allele-frequency in gnomAD) | 65 (6.2%) |
| Excluding longer than 10bp indels | 60 (5.8%) |
| Excluding adjacent indels within 10bp range | 58 (5.6%) |
| Excluding indels with Low-confidence score (LC) by LOFTEE | 46 (4.4%) |

**Supplementary Table 10**. **PLP variants with ≥1% frequency in the Dutch and the Estonian cohorts**
Shadowed lines represent variants with ≥1% carrier frequency in both cohorts; Frequencies >1% are marked in bold

| **Gene; Variant** | **Phenotype** | **Dutch cohort (N=4120)** | | | **GoNL Allele frequency (%) (N=498)** | **Estonian cohort (N=2327)** | | |
| --- | --- | --- | --- | --- | --- | --- | --- | --- |
|  |  | **Number of heterozygotes** | **Number of homozygotes** | **Allele frequency (%)** |  | **Number of heterozygotes** | **Number of homozygotes** | **Allele frequency (%)** |
| *HFE*; p.Cys282Tyr | Hemochromatosis | 441 | 18 | **11.58** | 5.4 | 176 | 3 | **7.82** |
| *BTD*; p.Asp444His | Biotinidase deficiency | 294 | 3 | **7.28** | 3.6 | 188 | 2 | **8.25** |
| *SERPINA1*; p.Glu288Val | Emphysema due to AAT deficiency | 264 | 5 | **6.65** | 3.9 | 75 | 1 | **3.31** |
| *SERPINA1*; p.Glu366Lys | Emphysema due to AAT deficiency | 139 | 3 | **3.52** | 1.7 | 75 | 1 | **3.31** |
| *RBM8A*;  c.-21G>A | Thrombocytopenia-absent radius syndrome | 93 | 3 | **2.40** | 2.6 | 63 | 0 | **2.71** |
| *CFTR*; p.Phe508del | Cystic fibrosis | 91 | 0 | **2.21** | 1.9 | 30 | 0 | **1.29** |
| *ACADM*; p.Lys362Glu | deficiency of Acyl-CoA dehydrogenase, medium chain | 82 | 0 | **1.99** | 0.6 | 3 | 0 | 0.13 |
| *OCA2*; p.Val419Ile | Albinism | 80 | 0 | **1.94** | 0.8 | 2 | 0 | 0.09 |
| *ABCA4*; p.Gly863Ala | Retinitis pigmentosa 19 | 78 | 0 | **1.89** | 0.8 | 6 | 0 | 0.26 |
| *ACY1*; p.Arg353Cys | Aminoacylase 1 deficiency | 67 | 0 | **1.63** | 0.3 | 24 | 0 | **1.03** |
| *GALC*; p.Thr112Ala | Krabbe disease | 54 | 0 | **1.31** | 1.0 | 5 | 0 | 0.21 |
| *PMM2*; p.Arg141His | Congenital disorder of glycosylation, type Ia | 50 | 0 | **1.21** | 0.8 | 9 | 0 | 0.39 |
| *TF*; p.Gly671Glu | Atransferrinemia | 45 | 1 | **1.14** | 0.5 | 16 | 0 | 0.69 |
| *CFTR*; p.Arg117His | Cystic fibrosis | 43 | 0 | **1.04** | 0.7 | 3 | 0 | 0.13 |
| *MUTYH*; p.Gly369Asp | Adenomas, multiple colorectal | 42 | 0 | **1.02** | 0.2 | 12 | 0 | 0.52 |
| *FGB*; p.Pro26q5Leu | Afibrinogenemia, congenital; | 42 | 0 | **1.02** | 0.7 | 4 | 0 | 0.52 |
|  | Hypofibrinogenemia, congenital |  |  |  |  |  |  |  |
| *GJB2*;  c.35del | Deafness, autosomal recessive 1A | 27 | 0 | 0.66 | 0.8 | 96 | 1 | **4.21** |
| *CLCN1*; p.Arg894Ter | Myotonia congenita, recessive | 17 | 0 | 0.41 | 0.0 | 75 | 0 | **3.22** |
| *LIPC*; p.Thr405Met | Hepatic lipase deficiency | 13 | 0 | 0.32 | 0.1 | 60 | 0 | **2.58** |
| *ALDOB*; p.Ala150Pro | Fructose intolerance, hereditary | 33 | 0 | 0.80 | 0.4 | 52 | 0 | **2.23** |
| *UGT1A1*; p.Gly71Arg | Crigler-Najjar syndrome, type I/II; Gilbert syndrome | 7 | 0 | 0.17 | 0.0 | 42 | 0 | **1.80** |
| *ABCA4*; p.Gly1961Glu | Retinitis pigmentosa 19 | 11 | 0 | 0.27 | 0.1 | 37 | 2 | **1.76** |
| *PAH*; p.Arg403Trp | Phenylketonuria | 1 | 0 | 0.02 | 0.0 | 36 | 0 | **1.55** |
| *NUP93*; p.Arg388Trp | Nephrotic syndrome, type 12 | 3 | 0 | 0.07 | 0.0 | 32 | 0 | **1.38** |
| *EDN3*; c.565dup | Waardenburg syndrome, type 4B | 12 | 0 | 0.29 | 0.1 | 31 | 0 | **1.33** |
| *TGM5*; p.Gly113Cys | Peeling skin syndrome 2 | 28 | 0 | 0.68 | 0.2 | 27 | 0 | **1.16** |
| *UPB1*;  c.917-1G>A | Beta-ureidopropionase deficiency | 15 | 0 | 0.36 | 0.2 | 27 | 0 | **1.16** |

**Supplementary Table 11. Diseases that are tested in the neonatal screening program with their abbreviations and gene names**

| **Disease** | **Abbreviation** | **Gene/s** |
| --- | --- | --- |
| Medium-chain acylCoA dehydrogenase deficiency | MCAD | *ACADM* |
| Very long chain acyl-CoA dehydrogenase deficiency | VLCAD | *ACADVL* |
| Maple syrup urine disease | MSUD | *BCKDHA/BCKDHB/DBT* |
| Biotinidase deficiency | BIO | *BTD* |
| Cystic-fibrosis | CF | *CFTR* |
| Tyrosinaemia type 1 | TYR-1 | *FAH* |
| Galactosemia | GAL | *GALT* |
| Glutaric aciduria type 1 | GA-1 | *GCDH* |
| Trifunctional Protein deficiency / Long-chain hydroxyacyl-CoA dehydrogenase deficiency | LCHAD | *HADHA/HADHB* |
| Isovaleric aciduria | IVA | *IVD* |
| 3-Methylcrotonyl-CoA carboxylase deficiency | 3-MCC | *MCCC1/MCCC2* |
| Phenylketonuria | PKU | *PAH* |

**Supplementary Table 12. P-values for permutations of consanguinity ratio (CR) based on coding length of the gene panels**

Significant scores are marked in Bold. Bonferroni correction was calculated for 13 tests.

|  | **Dutch cohort** | | | **Estonian cohort** | | |
| --- | --- | --- | --- | --- | --- | --- |
| **Panel** | **True CR** | **P-value two-tailed** | **After Bonferroni correction** | **True CR** | **P-value two-tailed** | **After Bonferroni correction** |
| **Skeletal** | 123.03 | **0.0004** | **0.01** | 91.87 | **0.02** | 0.29 |
| **ID** | 48.02 | **0.0004** | **0.01** | 40.86 | **0.003** | **0.04** |
| **Neuromuscular** | 47.28 | 0.10 | 1.00 | 9.53 | 1.00 | 1.00 |
| **Cilia+Kidney** | 36.72 | 0.52 | 1.00 | 18.16 | 1.00 | 1.00 |
| **Dermatologic** | 29.11 | 0.99 | 1.00 | 22.82 | 1.00 | 1.00 |
| **Immune system** | 22.62 | 1.00 | 1.00 | 28.71 | 0.86 | 1.00 |
| **Deafness** | 22.59 | 1.00 | 1.00 | 21.29 | 1.00 | 1.00 |
| **Hematologic** | 19.15 | 1.00 | 1.00 | 30.10 | 1.00 | 1.00 |
| **Multisystem disorders** | 18.69 | 0.92 | 1.00 | 12.97 | 1.00 | 1.00 |
| **Endocrine** | 14.38 | 1.00 | 1.00 | 13.40 | 1.00 | 1.00 |
| **Metabolic-ID** | 14.25 | 1.00 | 1.00 | 20.09 | 0.77 | 1.00 |
| **Metabolic** | 9.52 | 1.00 | 1.00 | 11.31 | 1.00 | 1.00 |
| **Blindness** | 7.12 | 1.00 | 1.00 | 6.58 | 1.00 | 1.00 |

**Supplementary Table 13. Data used for the analysis of selection signatures (excel)**

The table presents gene ID, gene transcript, chromosome, starting and ending coordinates, gene name, selection measurements scores and the gene-panel

**Supplementary Table 14. Variants classified as PLP in this study (excel).**

The table presents variant information for all variants that were classified as PLP in this study.
